# Pericytes enable effective angiogenesis in the presence of pro-inflammatory signals

**DOI:** 10.1101/608059

**Authors:** Tae-Yun Kang, Federico Bocci, Mohit Kumar Jolly, Herbert Levine, José Nelson Onuchic, Andre Levchenko

**Author notes:** To whom the manuscript related communications should be addressed (Herbert Levine, José Nelson Onuchic, Andre Levchenko).

## Abstract

Angiogenesis frequently occurs in the context of acute or persistent inflammation. The complex interplay of pro-inflammatory and pro-angiogenic cues is only partially understood. Using a new experimental model permitting exposure of developing blood vessel sprouts to multiple combinations of diverse biochemical stimuli and juxtacrine cell interactions, we present evidence that a pro-inflammatory cytokine, tumor necrosis factor (TNF), can have both pro- and anti-angiogenic effects, depending on the dose and the presence of pericytes. In particular, we find that pericytes can rescue and enhance angiogenesis in the presence of otherwise inhibitory high TNF doses. This sharp switch from pro- to anti-angiogenic effect of TNF observed with an escalating dose of this cytokine, as well as the effect of pericytes are explained by a mathematical model trained on the biochemical data. Furthermore, this model was predictive of the effects of diverse combinations of pro-and anti-inflammatory cues, and variable pericyte coverage. The mechanism supports the effect of TNF and pericytes as modulating signaling networks impinging in Notch signaling and specification of the Tip and Stalk phenotypes. This integrative analysis elucidates the plasticity of the angiogenic morphogenesis in the presence of diverse and potentially conflicting cues, with immediate implications for many physiological and pathological settings.

## Introduction

Developmental and physiological processes are frequently guided by diverse, sometimes conflicting cues. These cues may reflect a complex and evolving local environment, to which the process outcomes need to be matched. For instance, angiogenesis — the growth and morphogenesis of new vascular networks from existing ones — is triggered by the disruption of the local oxygen supply, encoded at the signaling level by a host of secreted factors^1, 2^. Angiogenesis also frequently occurs in the presence of pro-inflammatory stimuli, both acute, as in physiological wound healing, and persistent, as in the context of tumor growth and various pathologies, including asthma and chronic wounds, the local hypoxia is frequently accompanied by inflammatory conditions^3, 4^. Furthermore, these pro-inflammatory signals can have a direct effect on vascular stability, and the initiation and progression of angiogenesis^5-7^. The signaling cues in the local microenvironment can also change in time as the oxygen supply is gradually restored and as other concomitant processes, such as resolution of inflammatory response, unfold. Therefore, a proper control of vascular morphogenesis must involve a tight coordination of responses to diverse cues, including those involved in the immune response. However, in spite of decades-long research, it is not clear how this coordination is achieved and how it may be misregulated in various pathological settings, such as growing tumors, resulting in substantially altered structure and function of the vascular beds. Furthermore, the reported findings provide conflicting evidence as to whether the local inflammatory response may be pro- or anti-angiogenic^5, 7-9^. The complexity of the problem is further exacerbated by the intricate organization of vascular and immune systems, involving multiple cells types within highly organized cellular networks. Thus, we require new tools and approaches addressing the roles of distinct environmental and cell-generated cues in regulating angiogenesis or other complex morphogenetic processes.

Angiogenesis has been studied on diverse scales, from molecular networks to tissues, using a very diverse set of techniques. In particular, mimicking angiogenesis in various bio-engineered devices by multiple groups has allowed a careful untangling of the basic regulatory mechanisms governing this process^10-12^. Many of the inferred mechanisms have been confirmed *in vivo*, justifying the methodology, and leading to its use in tissue engineering efforts and medical interventions. However, a degree of simplification inherent in many of these methods frequently leaves important questions open. Therefore, a continued enhancement of the tissue modeling technologies is still needed to gain a better understanding of the complex underlying biology. Arguably, these developments can also benefit from computation modeling and theoretical analysis to account for salient features of complex intracellular and intercellular molecular interactions.

A key problem in the analysis of angiogenesis is how endothelial cells differentiate into diverse phenotypic states, including the Tip and Stalk cells^8, 9, 13, 14^. The Tip cells engage in locomotion within a hypoxic tissue leading the growing sprout, while Stalk cells undergo proliferation in coordination with the Tip cell locomotion to ensure the continuity of the growing sprout. These cells can undergo dynamic phenotypic switching, with some Stalk cells occasionally replacing the existing Tip cell or forming new Tip cells and spearheading new branches. Different phenotypic states can be induced by pro-angiogenic factors^15^ with the differentiation into distinct states further enhanced in neighboring cells by juxtacrine activation of the Notch signaling pathway^14^. As the emerging vessel matures, a lumen forms through a partially understood set of mechanisms, involving cell remodeling and reorganization^1, 16-18^. Both the phenotypic state (Tip vs. Stalk) selection and lumen formation can be strongly influenced by the relevant biochemically active ligands, including key pro-angiogenic growth factors, such as Vascular Endothelial Growth Factor (VEGF) and cytokines associated with inflammation, such as Tumor Necrosis Factor (TNF)^8, 9^. Whereas the effects of VEGF are thoroughly explored and well understood, the interplay between VEGF and TNF signaling and the resulting effects of these potentially conflicting cues remain a matter of debate, with little information available about the crosstalk of the pathways these ligands can trigger on the molecular level. Furthermore, although supportive mural cells, such as pericytes and smooth muscle cells, have been implicated in angiogenesis and maintenance of blood vessels^19, 20^, it is not clear how they might modulate the effects of VEGF and TNF, and other relevant cues on endothelial cells.

To address the need for a better, more quantitative understanding of the crosstalk between pro-angiogenic and pro-inflammatory stimuli, here we report on development of a new meso-scale fabrication technique allowing monitoring of angiogenesis on the single cell level in the presence of diverse sets of highly controlled cues and a highly controlled pre-patterning of endothelial cells and pericytes within a collagen matrix. We used this technique to study the effect of a large number of combinations of cues, in the presence and absence of co-patterned pericytes, on angiogenesis outcomes. To account for the experimental findings and to unravel the mechanisms controlling cell differentiation in response to diverse, potentially conflicting cues, we modified and extended a previously developed mathematical model of angiogenesis to account for the effects of TNF and pericytes^8^. We experimentally validated the model assumptions and predictions, and showed that it can account for various unexpected effects of the complex extracellular milieu. In particular, we demonstrate that the effect of TNF can be either pro- or anti-angiogenic, depending on its concentration and the other environmental inputs. We also show that pericytes can modulate the signaling processes activated by pro-angiogenic and pro-inflammatory cues, strongly modulating the phenotypic selection at the onset of angiogenesis and rescuing anti-angiogenic effects of TNF. These findings can assist in quantitative analysis and control of angiogenesis, particularly in the presence of the inflammatory response, in normal and pathological conditions.

## Methods

### Cell culture

Human umbilical vein endothelial cells (HUVECs) were purchased from Yale Vascular Biology Center and cultured in M199 (Gibco) supplemented with 20% FBS (Life Technologies), 1% HEPES (Thermo Scientific), 1% Glutamax (Thermo Fisher), 1% antibiotic-antimycotic (Thermo Fisher), Heparin (25mg/500ml, Sigma Aldrich), and endothelial cell growth supplement (Sigma Aldrich). Human pericytes (PCs) were kindly provide by Dr. Anjelica L. Gonzalez (Yale University, CT, USA). PCs were isolated by explant outgrown from microvessels from de-identified human placenta, characterized using previously established methods^21^, and cultured in M199 supplemented with 10% FBS, 1% antibiotic-antimycotic. As angiogenic and inflammatory factor, VEGF (100ng/ml, Gibco) and TNF-alpha (100ng/ml, Gibco) were used, respectively. For inducing prolonged and matured vessel formation in 3D vessel chip, 40ng/ml of basic fibroblast growth factor (bFGF, Thermo Fisher), 500nM of Sphingosine-1-phosphate (S1P, Sigma Aldrich) and 75ng/ml of phorbol myristate acetate (PMA, Sigma Aldrich) were mixed along with 100ng/ml of VEGF.

### Biomimetic 3D Vessel fabrication

The biomimetic 3D vessel chip consists of a PDMS chamber, an engineered blood vessel embedded in collagen gel and a cover slip and they were assembled without external jigs. All templates for PDMS and collagen casting were designed by SolidWorks and the cad files were converted to gcode for a commercialized 3D printer (Ultimaker) through Cura (Ultimaker). PLA filament was used for building structures. The details of the fabrication process is described in Fig. S1. Once HUVECs formed a confluent monolayer on the channel surface, fresh medium was injected through the inlet hole and medium was changed every 6 hours before the treatment with pro- or anti-angiogenic factors.

### Co-culture of endothelial cells and pericytes

Endothelial cells and pericytes were cultured with transwell inserts for 6-well plates as described in Fig. 3A. To co-culture endothelial cells and pericytes in a layered configuration, both sides of a insert membrane (Corning), which has 24mm diameter, 0.4 µm pore size, and 10 um thickness, were coated with laminin (10 µg/ml). Pericytes were first seeded on the bottom side of the membrane and allowed to adhere for 2 hours. Then the transwell insert was placed in a 6-well plate and endothelial cells were seeded on the top side of the membrane. After overnight incubation, TNFα and VEGF were added on both apical and basal sides.

### Sandwich culture

A 24-well plate was coated with 0.2 ml of collagen mixture (5mg/ml) for each well and incubated for 1 hour at 37 °C to form a collagen gel. After gelation, endothelial cells with or without pericytes were seeded on the gel and incubated at 37 °C overnight. The total cell number on a gel was set to be 2 × 10^5^ and the ratio of endothelial cells and pericytes was 9:1. On the following day, the culture medium was gently removed and 0.2 ml of collagen mixture was added again and incubated for 1 hour at 37 °C. After gelation, 1 ml of medium was applied with TNFα or VEGF.

### Western blot

Endothelial cells were collected with plastic scrapers carefully without damaging the membrane after PBS washing. Western blotting was used for measuring protein expression levels. Cells were lysed with lysis buffer (RIPA buffer) following the manufacturer’s protocol. The cell lysates were heated with Laemmli buffer at 95 °C for 5 min. Then the lysates were loaded to 4-20 % gels (Bio-Rad) for electrophoresis and proteins were transferred to a nitrocellulose membrane. The membrane was blocked with 3% BSA in TBST and incubated with primary antibodies overnight at 4 °C and followed by horseradish-peroxidase-coupled secondary antibody for 1 hour at room temperature. Between each step, the membrane was washed three times with TBST for 15min each. Finally, the membrane was incubated with ECL for revealing bands through ChemiDoc XRS (Bio-rad). The bands were quantified with ImageJ and normalized to GAPDH expression. All primary antibodies were used at 1:1000 dilution and secondary antibodies were used at 1:2000 dilution. Antibodies used in western blotting are as follows: Jagged-1 (cell signaling), Dll4 (cell signaling), pNFkB (cell signaling), NFkB (cell signaling), pErk (cell signaling), Erk (cell signaling), pJNK (cell signaling), JNK (cell signaling), GAPDH (cell signaling), HRP-linked anti-rabbit (GE HealthCare), HRP-linked anti-mouse (GE HealthCare),

### Immunofluorescence staining

For all immunofluorescence staining, cells were fixed with 4% (wt/wt) formaldehyde for 20 min, permeabilized with 0.1% triton-X for 10 min, and blocked in 10% goat serum for 1hour. All primary antibodies were used at 1:50 dilution and secondary antibodies were used at 1:100 dilution. The primary and secondary antibodies used in immunofluorescence staining are as follows: VE-Cad (Santa Cruz), α-SMA (Abcam), anti-mouse 488 (Thermo Scientific), anti-rabbit 594 (Thermo Scientific). Hoechst and and Alexa Fluor 594 phalloidin were used at 1:500 dilution and 1/200 dilution, respectively. Pre-labeling of cells were performed with Vybrant cell-labeling solution (Thermo Fisher) following the manufacturer’s protocol.

### Image acquisition and analysis

Sandwich culture images were acquired with 20× objective attached to phase contrast microscopy. The resulting tube-like networks were quantified with Angiogenesis Analyzer for ImageJ. Confocal immunofluorescence images were acquired with 20× water-immersion objective attached to Leica scanning disk. Either Leica software or IMARIS was used to merge channels, stack layers for 3D reconstruction and generate longitudinal and transverse cross-sections. IMARIS was used to count the numbers of sprouting and to measure the length of newly formed sprouts. At least two samples were prepared for each condition and the quantified data were combined together.

### Statistical analysis

Sample populations were compared using one-way ANOVA. P < 0.05 was the threshold for statistical significance.

### Mathematical model

To unravel the interplay between Notch, VEGF and TNF signaling pathways, we extended the mathematical model describing the interaction of Notch and VEGF developed by Boareto et al^8^ to the Notch-VEGF-TNF signaling axis. The equations and parameters are discussed in supporting information. The computational analysis was performed using the Python numerical library PyDSTool^22^. The model construction and parameters used for the model are discussed in supplementary information.

## Results

To analyze the effects of different combinations of molecular pro-angiogenic and pro-inflammatory cues and of mural cells on angiogenesis, we engineered a new biomimetic system allowing precise structuring of an artificial blood vessel, with controlled juxtaposition of layers of endothelial cells and pericytes within a collagen gel (Figs. 1A-C & Fig. S1, see Methods for details). We next demonstrated that this experimental system devoid of non-biological materials enabled generation of a basement membrane separating polarized endothelial cells and abluminal pericytes, with pericyte assuming elongated morphology with processes, characteristic of the *in vivo* endothelial coverage (Fig. 1D). We note that, unlike other previously described blood vessel models with mural cells^23-26^, the major advantage of this newly developed experimental system is to enable a highly controlled juxtaposition and possible interaction between endothelial cells and pericytes from the outset of the experiment, without any additional treatment used to recruit pericytes onto endothelium from the surrounding gel. The cells were accessible to high resolution imaging modalities, including confocal microscopy, which was used throughout the experimental analysis presented here. Functionally, one of the effects of pericytes is to increase the micro-vessel stability, which is reflected in lower leakage and permeability to luminal substances. Indeed, we found that the vascular permeability of lumenal FITC-dextran was decreased by 40% in the presence of pericytes vs. the pericyte-free version of the system (Figs. S2A-D).

**Figure 1.**
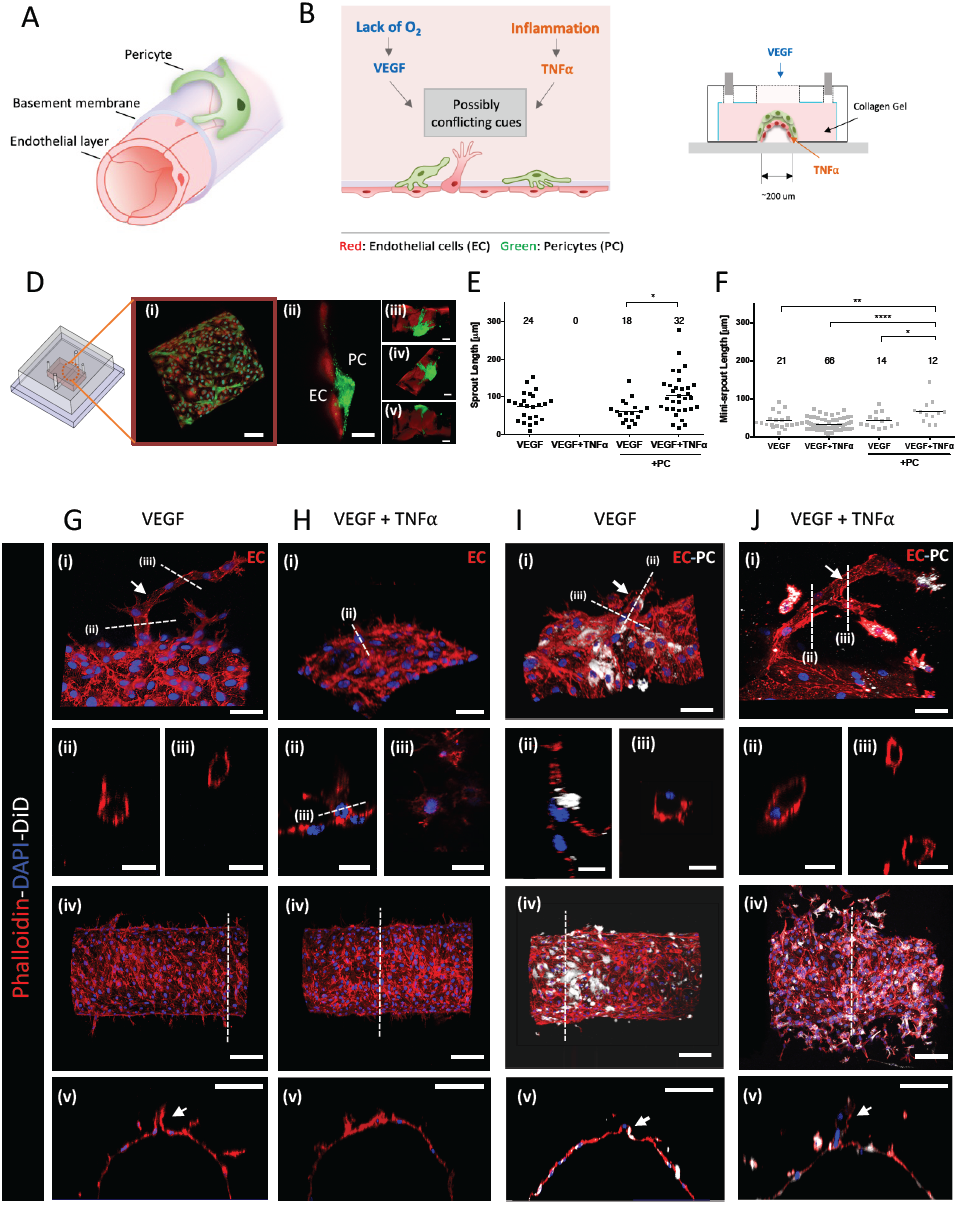
3D vessel on a chip effectuating the interaction between endothelium and pericytes. (A) Layered structure of a capillary vessel: endothelium, basement membrane, and pericytes. (B) Schematic description of angiogenesis co-stimulated by cues from inflammation and ischemic tissue. This study focuses on how those possibly conflicting cues control angiogenesis in multi-cellular vessel structure. (C) Schematic diagram of 3D vessel on a chip mimicking the organization of capillary vessel embedded in collagen type I. (D) Confocal images of endothelial cells (red) and pericytes (green): (i-ii) Endothelial cells formed a monolayer on the channel and pericytes were covering the endothelium in close proximity. Scale bar, 100 *μm*. (iii-v) Basal sides of endothelial cells and pericytes were facing each other. Scale bar, 20 *μm*. Quantified lumenized sprouts (E) and single cell-sized mini-sprouts (F) from angiogenesis assay on the 3D vessel on a chip. Confocal images of lumenized sprouts formation in response to gradient of VEGF (G), single cell-sized mini-sprouts formation in response to gradient of VEGF and local TNFα (H), shorter sprouts in response to gradient of VEGF in the presence of pericyte (I), the rescued sprout formation in response to gradient of VEGF and local TNFα in the presence of pericytes (J). Actin filaments of endothelial cells and pericytes were stained with phalloidin (red) and pericytes were pre-labeled with DiD (white). White arrows in (G, ii-iii), (I, ii-iii) and (J, ii-iii) indicate sprouts with hollow lumens. Scale bar, 50 *μm* in (i), 25 *μm* in (ii) & (iii), 100 *μm* in (iv) & (v) of G-J. Cocktail medium was used as a basal medium for all samples.

To further investigate the utility of this experimental system for the analysis of angiogenesis, we supplied exogenous pro-angiogenic and pro-inflammatory cues (Fig. 1C). In particular, we delivered a previously reported^11^ pro-angiogenic cocktail by allowing it to diffuse the collagen gel surrounding the engineered vessel. This cocktail contained VEGF supplemented with 40 ng/ml of basic fibroblast growth factor (bFGF), 500 nM of sphingosine-1-phosphate (S1P) and 75 ng/ml of phorbol 12-myristate 13-acetate (PMA). We found that, with or without pericytes, within three days, this cocktail indeed triggered extensive angiogenesis resulting in multiple lumenized sprouts, extending many cell diameters from the parental vessel, frequently with multiple branches (Figs. 1E-J). Notably, if 100 ng/ml of TNF was locally supplied in addition to the pro-angiogenic cocktail, in the absence of pericytes, the formation of long, lumenized sprouts was completely abolished (Figs. 1E,H). Instead, we observed the formation of single cell-sized mini-sprouts, protruding from the ablumenal side of the parental vessel wall (Fig. 1F). This response suggested that 100 ng/ml of TNF had an essentially anti-angiogenic effect, perturbing a key aspect of successful lumenal sprout formation. Strikingly, this anti-angiogenic effect of TNF was completely rescued if the experiment was repeated in the presence of pericytes covering the abluminal side of the engineered vessel (Figs. 1E,J). Surprisingly, we also found that many of the sprouts forming under these conditions were longer vs. those observed in the absence of TNF, suggesting an additional pro-angiogenic effect of the TNF-pericyte combination. On the other hand, the effect of pericytes on VEGF-mediated angiogenesis in the absence of TNF was relatively minor, with slight decrease of the number but not the length of the sprouts (Figs. 1E,I). Overall, these results supported the experimental assay as a controllable model of angiogenesis. More importantly, our findings revealed a complex control of angiogenesis by multiple cues presented in different combinations, with the effect of TNF strongly modulated by the presence of pericytes.

Due to the complex nature of the pro-angiogenenic cocktail, we explored if VEGF signaling alone might interact with the cues provided by TNF and pericytes in a fashion similar to that of the whole cocktail (Figs. 2A-F). We found that, in the absence of pericytes, 100 ng/ml of VEGF alone induced a very limited but measurable effect, promoting the formation of single-cell mini-sprouts (Fig. 2A), but not the lumenized longer sprouts enabled by the full cocktail. Interestingly, however, we observed that even this limited effect of VEGF was completely abrogated if 100 ng/ml of TNF was also supplied to the cell environment (Fig. 2C). This result was again suggestive of anti-angiogenic effect of TNF, even in the context of a limited pro-angiogenic phenotype promoted by VEGF. We found that the effect of TNF was partially reversed in the presence of pericytes, leading to a more limited formation of mini-sprouts (Fig. 2D) vs. the effect of VEGF in the absence of pericytes and TNF (approximately two-fold lower mini-sprout formation) (Fig. 2B). The experiments also suggested that pericytes without TNF had a more pronounced inhibitory effect on VEGF-induced mini-sprout formation (number of mini-sprouts) vs. either sprout or mini-sprout formation induced by the full cocktail (Figs. 2E,F). Overall, these results supported the rescue effect that pericytes can have on the anti-angiogenic TNF signaling, even in the presence of a weak pro-angiogenic signal leading to incomplete sprout formation.

**Figure 2.**
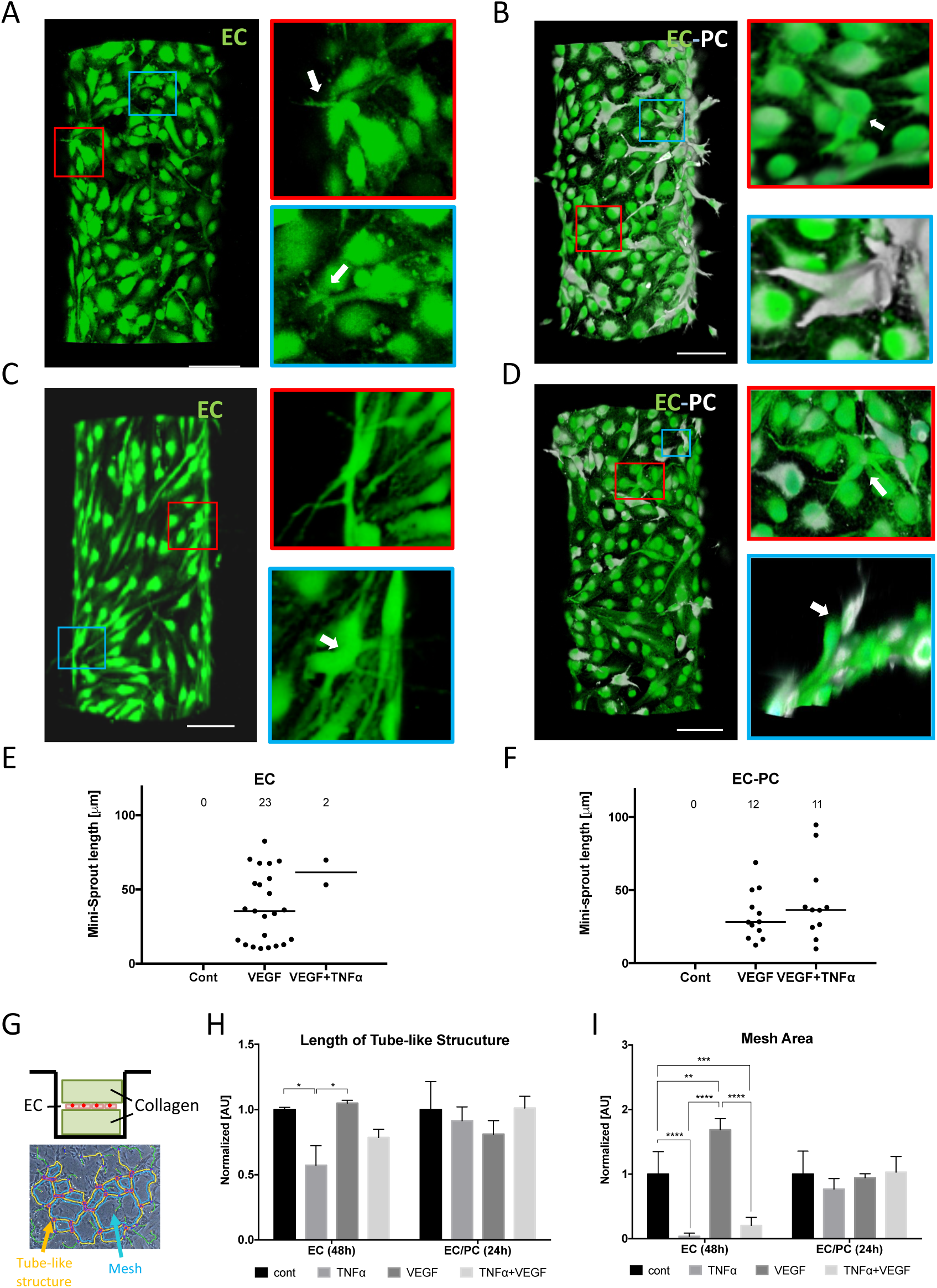
Interplay of TNFa and VEGF on initiation of angiogenesis and its alteration by pericytes. Representative images of mini-sprouts formation and cell migration induced by VEGF without (A) and with (B) pericytes. Inhibited mini-sprouts formation by TNF*α* without pericytes (C) and rescued mini-sprouts formation by pericytes (D). GFP-tagged endothelial cells (green) and DiD pre-labeled pericytes (white). Scale bar, 100 *μm*. White arrows indicate single-cell mini-sprouts which are the limited angiogenic response in normal growth medium without any supplement. Quantification of mini-sprouts formation without (E) and with (F) pericytes. (G) Experimental setup for vasculogenesis assay. Endothelial monolayer was cultured between two collagen layers and TNF*α* and VEGF were added with normal culture medium. Total length of tube-like structure (H) and mesh area (I) were significantly inhibited by TNF*α* on EC monolayer, while the difference was diminished in EC/PC mixed monolayer.

We also performed a more traditional vaculogenesis-mimicking assay relying on culturing cells being ‘sandwiched’ between two slabs of collagen gel (Fig. 2G). As expected, this cell culture method resulted in the characteristic mesh networks indicative of endothelial cell self-organization, thought to be reflective of conditions also leading to the vascular bed formation *in vivo* (Figs. 2G, S3A-H). We again found that the formation of these networks was significantly perturbed by TNF, but the inhibitory effect was rescued by the presence of pericytes (Figs. 2H & I).

Given the observed effects of TNF and pericytes, we next investigated the molecular mechanisms of crosstalk between pro- and anti-angiogenenic cues and their modulation by pericytes. Successful angiogenesis relies on a coordinated differentiation of endothelial cells in the parental vessel into the Tip and Stalk phenotypic states. This process is regulated by activation of the Notch mediated signaling, serving, as in many other differentiation contexts, to induce different cell fates in adjacent cells. Two Notch-ligands have been strongly implicated in regulation of Notch activity during angiogenesis: Delta-4 (Dll4) and Jagged-1 (Jag1)^8, 9, 14^. Furthermore, VEGF and TNF can induce the expression of Dll4 and Jag1, respectively^9, 27^. We therefore investigated how the signaling networks specific to VEGF and TNF might interact with each other in the presence or absence of pericytes. To enable this analysis, we simplified the experimental system by culturing ECs as a monolayer, both in isolation and in co-culture with pericytes (Fig. 3A). Two types of co-culture were used. In the first, the pericytes were cultured on a porous membrane within an insert that was within the same culture medium as the endothelial cells, enabling spatial separation of the heterotypic cells, but also a possibility of paracrine interactions between them (EC/PC co-culture). In the second, the endothelial cells and pericytes were co-cultured on two sides of the same porous membrane, which enabled both contact-mediated and paracrine interactions (EC-PC co-culture). Indeed, we observed endothelial cells and pericytes making contact through the 0.4 micron pores and forming N-cadherin-rich junctions in EC-PC co-culture (Fig. 3B). The results in the above co-culture experiments were contrasted with those from an endothelial monolayer monoculture experiments (EC culture), used as a control condition.

**Figure 3.**
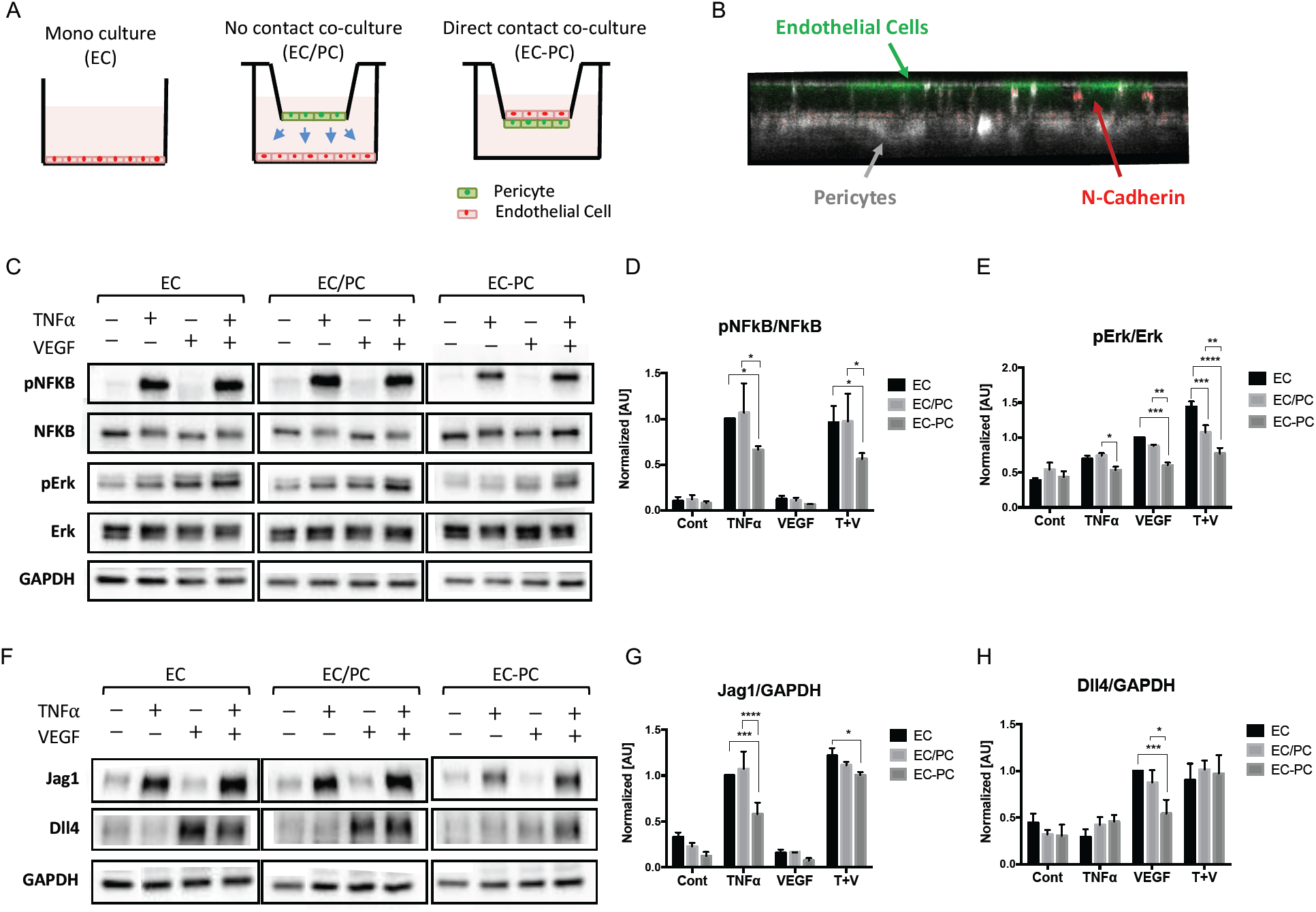
Inhibition TNFa and VEGF mediated signaling pathways on endothelium in a pericyte contact-dependent fashion. (A) Experimental setup for mono culture (EC), no contact co-culture (EC/PC), and direct contact co-culture (EC-PC) of ECs and PCs using transwell inserts. (B) ECs (green) and PCs (white) sitting across a permeable membrane. ECs and PCs are making N-cadherin (red) adhesion through pores of the transwell membrane. (C) Phosphorylation levels of TNF*α* and VEGF downstream targets analyzed by western blots. The bar graphs of pNFkB/NFkB (D) and pErk/Erk (E) were quantified from blot images (n=3). (F) Notch ligand Jag1 and Dll4 expression analyzed by western blots. The bar graphs of Jag1/GAPDH (G) and Dll4/GAPDH (H) were quantified from the blot images (n=3). Pericytes downregulated the downstream of VEGF and TNF in endothelial cells, in a cell contact-dependent fashion.

We then evaluated the signaling responses to TNF or VEGF alone, or the mixture of these signals. In particular, we analyzed the phosphorylation of NF-kappaB and Erk at 10 min. following the stimulation (the time at which these pathways display a high acute activation), as these pathways are both implicated in specific responses to these ligands, and as mediating Jag1 and Dll4, respectively. We found that NF-kappaB was indeed potently activated by TNF, whereas VEGF primarily activated Erk, under all conditions (Figs 3C-E, Figs. S4A & B). Furthermore, the effect of the combination of VEGF and TNF was additive, suggesting a limited crosstalk between the signaling pathways downstream of the receptors. We then investigated the effect of pericytes on the signaling outcomes. In the EC/PC co-culture, we found that pericytes had no significant effect on the phosphorylation of the signaling molecules (Figs. 3C-E), with a possible exception of JNK (Figs. S4A & B). On the other hand, in the EC-PC co-culture, there was a strong and highly significant inhibitory effect on all signaling molecules (Figs. 3C-E). These results suggested that pericytes inhibit signaling by both VEGF and TNF in a contact-dependent fashion.

We then explored if these short-term signaling effects could be reflected in the longer term changes in the expression levels of Jag1 and Dll4. We found, as expected, that, at 16 hrs. following exposure to signaling inputs, VEGF specifically induced an increased the expression of Dll4, whereas TNF enhanced the expression level of Jag1(Figs. 3F-H). We further observed that a combination of these ligands induced the expression of Dll4 and Jag1 to levels induced by each of the corresponding signaling inputs alone, again suggesting a very limited crosstalk between the signaling pathways. We also again did not observe a significant effect of pericytes on expression of these two molecules in the EC/PC co-culture (Figs. 3F-H). However, in the EC-PC co-culture case, we found a significant downregulation of Jag1 and Dll4 in response to TNF and VEGF inputs respectively (Figs. 3F-H). Interestingly, in spite of the additive signaling effect of the two ligands, the influence of pericytes on the combined action of TNF and VEGF in the EC-PC co-culture was much more muted, but has nevertheless resulted in a significant reduction of Jag1 levels vs. the control case (Figs. 3F-H). Overall, these results suggested that pericytes may downregulate the VEGF and TNF induced expression of Dll4 and Jag1 in endothelial cells, in a cell contact-dependent fashion, in a manner consistent with the effects of pericytes on the signaling pathways triggered by these pro- and anti-angiogenic factors. However, these results also raised the question of why the effect of pericytes in the EC-PC co-culture on expression of Dll4 was not significant. More generally, it was not clear how these molecular interactions might quantitatively translate into angiogenic outcomes. We thus next sought an explanation to these questions and the angiogenic response more generally through a combination of mathematical modeling and computational analysis.

To further investigate the possible interplay between TNF and VEGF signaling in the presence of pericytes and its effect on the expression of Jag1 and Dll4, and, ultimately, angiogenesis, we developed a mathematical model, partially based on a previous study^8^. Fig. 4A shows the schematic diagram describing the interactions between Notch, VEGF and TNF signaling. VEGF and TNF are treated as extracellular input signals, whereas the intracellular signaling networks consist of three interconnected modules: the Notch module, the VEGF response module and the TNF response module. The Notch module models signaling due to the engagement of the Notch receptor by the Dll4 and Jag1 ligands, transduced by the cleaved Notch Intracellular Domain (NICD). The VEGF and TNF modules describe lumped signaling pathway activations in response to each of these ligands. These modules interact with the Notch module through a crosstalk mechanism as described below^27^. First, TNF induces the expression of Jag1 by activating NF-kappaB. Moreover, NCID can transcriptionally inhibit the expression of the VEGF receptor (VEGFR2)^28, 29^, whereas activated VEGF module (AVEGF) induces the expression of Dll4. Finally, according to the published studies^29^, and as captured by the previous model^8^, activated Notch module can suppress Dll4 transcription and induce Jag1 transcription in the same cell. These interactions have been modeled as a series of ordinary differential equations, as described more in detail in the Supporting Information. Finally, we also introduced the effect of pericytes into the model by specifying the degree to which these cells can suppress the activation of Dll4 and Jag1 (Fig. 4B), based on the experimental data above (Fig. 3). Since the key phenotypic outcome regulated by these signaling pathways and controlling the initiation and progression of angiogenesis is the specification of the Tip and Stalk cells in adjacent cells, we further implemented the model on the scale of two adjacent cells, focusing, in particular, on whether the signaling interactions would result in differentiation of these model cells into the divergent phenotypic states.

**Figure 4.**
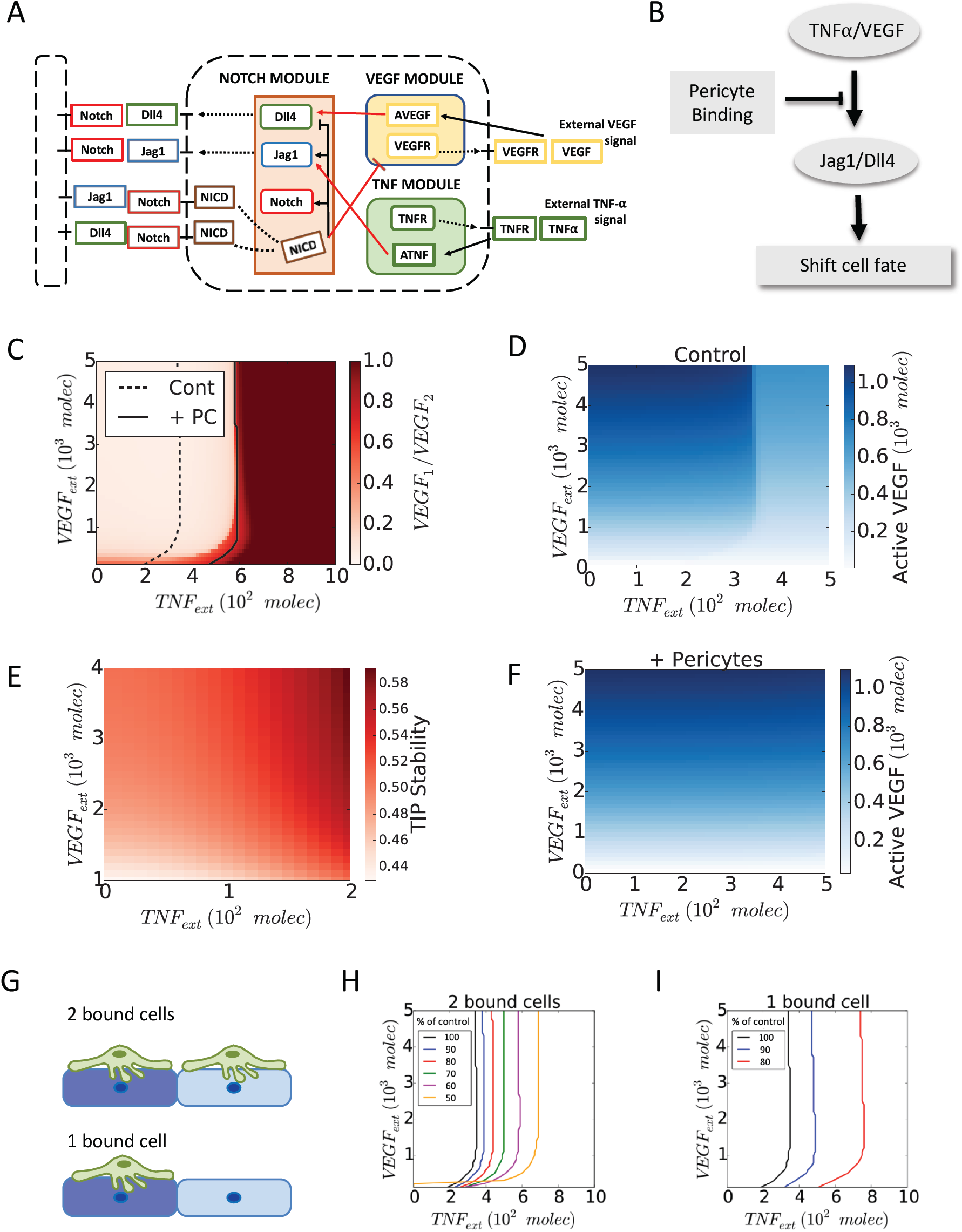
Pericytes shift the transition line between two distinct tip-stalk fate decision phases of endothelial cells. (A) Schematic diagram of the connections between Notch, VEGF and TNF*α* signaling. Red arrows highlight the crosstalk between the three components. Notch Intra Cellular Domain (NICD) transcriptionally inhibits VEGF receptor (VEGFR), while activated VEGF signaling (AVEGF) induces Dll4). Similarly, activated TNF*α* signaling (ATNF) induces Jagg1. (B) Schematic summary of the role of pericyte binding on notch signaling pathway and cell fate decision in angiogenesis. (C) Phase diagram of the relative VEGF activity in a 2-cell system for different levels of external TNF*α* (x-axis, *TNF*_*ext*_) and VEGF (y-axis, *VEGF*_*ext*_) signal. Pericytes slightly modify the action of external VEGF while inhibiting completely the effect of external TNF*α*. (D) Heat map of the level of active VEGF signaling in one of the two cells in bare endothelium. (E) The stability of the Tip cells by of external TNF*α* and VEGF (see Fig. S8 for details on calculation). (F) Heat map of the level of active VEGF signaling in one of the two cells in the presence of pericytes. (G) Description of 2 bound cells system and 1 bound cell system. Green cells and blue cells indicate pericytes and endothelial cells, respectively. Variation of the transition line upon the strength of inhibitory effect of pericytes in 2 bound cells system (H) and 1 bound cell system (I). As the value of p decreases, the transition line shifts further toward higher external TNF*α* signal. The value of p represents the percentage of Dll4 and Jag1 expression reduced by pericytes.

Analysis of the model revealed that cell exposure to combinations of VEGF and TNF concentrations result in diverse phenotypic responses, including the Tip phenotype (classified according to high levels of VEGFR and Dll4), the Stalk phenotype (low VEGFR, Dll4), as well as a less frequently discussed hybrid Tip/Stalk phenotype with intermediate levels of VEGFR, Dll4 (Supplementary figures S^5^, see details in the Supplementary). These results were summarized as a ‘phenotypic phase diagram’ (Fig. 4C). This diagram revealed that the degree of cellular differentiation expressed as the ratio of VEGFR activation levels in two adjacent cells increased gradually with the increasing VEGF input. This outcome was mediated by a gradual change in VEGFR activity in each of the cells (Fig. 4D). Interestingly, the model also predicted a putatively pro-angiogenic role of TNF, through promoting the Tip fate outcome, up to the threshold level (Fig. 4E). When the TNF dose exceeded this threshold level, there was a very abrupt abrogation of the cell differentiation, leading to an undifferentiated (or ‘hybrid Tip/Stalk’) state corresponding to similar levels of VEGFR activation in both of the modeled cells. This undifferentiated state of two neighboring cells was expected to disrupt effective angiogenesis, although individual cells might still adopt phenotypes promoting migration or differentiation, as described more in detail below. Overall, the model predicted that TNF-induced Notch-Jag1 signaling initially stabilizes a Tip phenotype putatively leading to pro-angiogenic responses at low doses, but prevented Notch-Dll4-driven Tip-Stalk differentiation, and thus was anti-angiogenic at higher doses, exceeding a sharp threshold. These findings were consistent with the experimentally observed anti-angiogenic effect of TNF at high doses (Figs. 1E-J, 2A-F), but left open the question of whether lower doses of TNF, below the predicted threshold, would indeed be pro-angiogenic, as predicted by the model.

Finally, we investigated the effect of pericytes on the modeling outcomes. In the case of dense coverage, allowing both model cells to receive the pericyte input (2 bound cells in Fig. 4G), we found that, for experimentally defined parameters, pericytes could completely rescue the anti-angiogenic effect of high TNF doses (Fig. 4C,F). Since pericytes make structurally complex contacts with endothelial cells, even the complete coverage might result in differential degree of pericyte input. We thus explored the graded effect of pericytes, finding that it resulted in a progressive shift of the threshold boundary formed by the critical TNF inputs for different VEGF levels (Fig. 4G, H). Strikingly, if the coverage was incomplete, so that only one cell in the cell pair was in contact with the pericyte (1 bound cell in Fig. 4G), the effect of graded pericyte input was much more pronounced, with only a moderate change of this input leading to a strong shift of the TNF threshold and thus more pronounced pericyte-mediated rescue effect (Fig. 4I). Indeed, Notch-driven cell differentiation emerges from dynamical competition among neighbors, and partial pericyte coverage could potentially provide an additional bias to cell-fate decision^30^. We then sought to validate these predictions in the experimental setting modeling angiogenesis in various defined combination of VEGF and TNF ligands described in Fig. 1.

One of the model predictions is that TNF, by inducing Jag1, indirectly suppresses Dll4 expression, consistent with prior literature reports^9^, which may also account for its negative effect on Tip vs. Stalk differentiation beyond a threshold level. This prediction also suggests that a combination of VEGF and TNF would exert positive and negative effects on Dll4 expression, respectively. Therefore, although both VEGF and TNF signaling may be attenuated by pericytes, when these molecular factors are present simultaneously, their attenuations can essentially cancel each other and thus pericyte input might not strongly affect Dll4 expression, as indeed observed above (Figs. 4C-E, S8A-C). On the other hand, the negative pericyte effect on the expression of Jag1 would be specific to attenuation of TNF signaling only and thus would still be predicted to occur under the co-stimulation conditions, again in agreement with the experimental data (Figs. 1E-J, 2A-F).

A more stringent test of the model can be provided by experimental exploration of the model space represented in the ‘phase diagram’ shown in Fig. 4C. To accomplish this, we exposed the cells in the 3D *angiogenesis assay described in Fig*. *1* to six combinations of different concentrations of VEGF (in the presence of the pro-angiogenic cocktail components) and TNF, in the presence or absence of pericytes (Table 1). To provide a quantitative assessment of the angiogenic response (Fig. S9), we evaluated several parameters of the emerging sprouts. They included the number of branches, the sprout length (in the case of several branches, we recorded the longest distance from a branch tip of the sprout to the root of the sprout connecting it to the parental vessel), the lumenized and multicellular nature of the sprouts, as well as the number of cells which have been assigned the Tip and Stalk fates (see the Methods section for details of this analysis). Using these metrics, we found the followings.

**Table 1.**
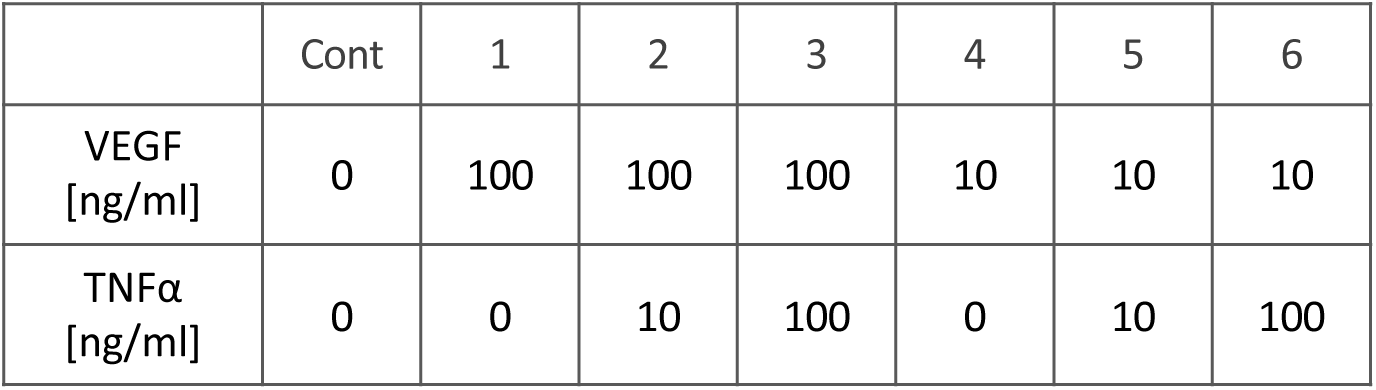
TNFα and VEGF concentration used for angiogenesis assay in 3D vessel on a chip.

|  | Cont | 1 | 2 | 3 | 4 | 5 | 6 |
| --- | --- | --- | --- | --- | --- | --- | --- |
| VEGF<br>[ng/ml] | 0 | 100 | 100 | 100 | 10 | 10 | 10 |
| TNFα<br>[ng/ml] | 0 | 0 | 10 | 100 | 0 | 10 | 100 |

Without TNF and in the absence of pericytes, in agreement with the model, the angiogenic response increased with increasing VEGF dose, yielding a greater number of lumenized sprouts, which were longer and more branched (points 1 vs. 4 in Figs. 5A,B). Also, in agreement with the model predictions, at each VEGF dose, escalating TNF levels were initially pro-angiogenic, increasing the number and branching of sprouts, as well the Tip cell formation (cf. points 1,2,3 and 4,5 and Fig. 5E). However, also agreeing with the model prediction, there was a complete abrogation of the formation of lumenized sprouts beyond a critical TNF level. The strikingly abrupt nature of this threshold phenomenon was underscored when cells were exposed to the combination of 100 ng/ml TNF and 100 ng/ml VEGF (the maximal amounts for both ligands used here). This was the only combination of inputs that provided two types of outcomes, varying from experiment to experiment. In two out of six independent experiments (Fig. 5C, Fig. S10A), there was an extensive formation of highly branched sprouts (point 3 in Fig. 5A), whereas in four out of six independent experiments (Fig. 5C & Fig. S10A), the lumenized sprout formation was completely shut down (denoted as point 3’ in Fig. 5A). This result indicated that this combination of TNF and VEGF inputs led to a divergent outcome due to being very close to the very sharp threshold separating the pro- and anti-angiogenic effects of TNF, with the outcome defined by a slight variation of experimental conditions. This allowed us to connect this experimental point with the phenotypic phase diagrams shown in Figs. 4C,D), indicating the position of the threshold levels. The model also predicted that the threshold TNF levels would shift slightly to the lower concentration values for lower VEGF inputs (Fig. 4C,D). In agreement with this prediction, when the VEGF concentration was lowered to 10 ng/ml, while keeping TNF levels at 100 ng/ml (point 6 in Fig. 5A), we observed an unambiguous and complete inhibition of lumenized sprouting in all experimental repeats (Fig. 5B). We noted that, at the TNF levels exceeding the threshold, there was still formation of single-cell mini-sprouts (Fig. 5D) and Tip cells (Fig. 5E), which however, were not supported by sprout growth through cell division and thus formation of Stalk cells (Fig. 5E), thus not resulting in functional angiogenesis. Finally, we found that, phenotypically, the application of VEGF without other components of the pro-angiogenic cocktail with or without TNF (as initially analyzed in Fig. 2), was equivalent to the responses to the full pro-angiogenic cocktails with 10-fold lower VEGF contents (gray dots in Fig. 5D, and open symbols adjacent to points 4 and 6 in the diagram in Fig. 5A), suggesting that the components of the cocktail act through the same molecular mechanisms in inducing angiogenic responses as those classically attributed to VEGF inputs, enhancing the VEGF signaling beyond what may be saturating levels.

We then experimentally examined the quantitative characteristics of the rescue effect of pericytes on angiogenesis, at TNF levels above the threshold values and different VEGF concentrations (+PC in Figs. 5B,D,E). For both VEGF concentrations, at 100 ng/ml of TNF (points 3’ and 6 in Fig. 5A), we observed a complete rescue of the angiogenesis, in a VEGF dependent fashion, which was consistent with the model predictions. Strikingly, at the higher VEGF input (points 1 and 3’ in Fig. 5A), the effect of pericytes was not only to rescue but to enhance the angiogenesis, leading to sprouts that were much longer (data indicated as +PC:1 and +PC:3 in Figs. 5B,D,E) than those observed for any of the VEGF/TNF combinations, in the absence of pericytes. This effect was consistent with an increased formation of Stalk cells when pericytes were present along with application of the highest levels of TNF and VEGF (Fig. 5E). A closer inspection of these sprouts revealed a high degree of variability in length and in the cell density (which was also associated with the sprout thickness) (Fig. 5F). We therefore explored if this effect might be related to a variable pericyte coverage at the Tip/Stalk cell area during the sprout extension. We found that, for the condition shown as point 3 in Fig. 5A, a fraction of the sprouts was associated with a single pericyte cell at the sprout tip area. These sprouts were on average significantly longer and less dense (thinner) than the sprouts forming without a pericyte cell at the tip (Figs. 5H). However, pericytes at the sprout tip area were only very rarely observed in the absence of TNF (Fig. 5G), resulting in a more variable sprout length and cell density distributions. These data suggested that the presence of a partial pericyte coverage at the threshold combination of VEGF and TNF (points 3,3’ of Fig. 5A) can substantially improve the stability of the differentiation process, leading to more persistent migration and division phenotypes and thus longer sprouts. These data again supported the mechanism proposed here to explain the angiogenesis outcome at diverse combinations of TNF and VEGF inputs, validating the predictions of the associated mathematical model. Overall, our results suggested that a combination of diverse inputs of pro- and anti-angiogenic stimuli and pericyte coverage can strongly modulate the outcome of angiogenesis, resulting in wide range of shapes of incipient sprouts.

**Figure 5.**
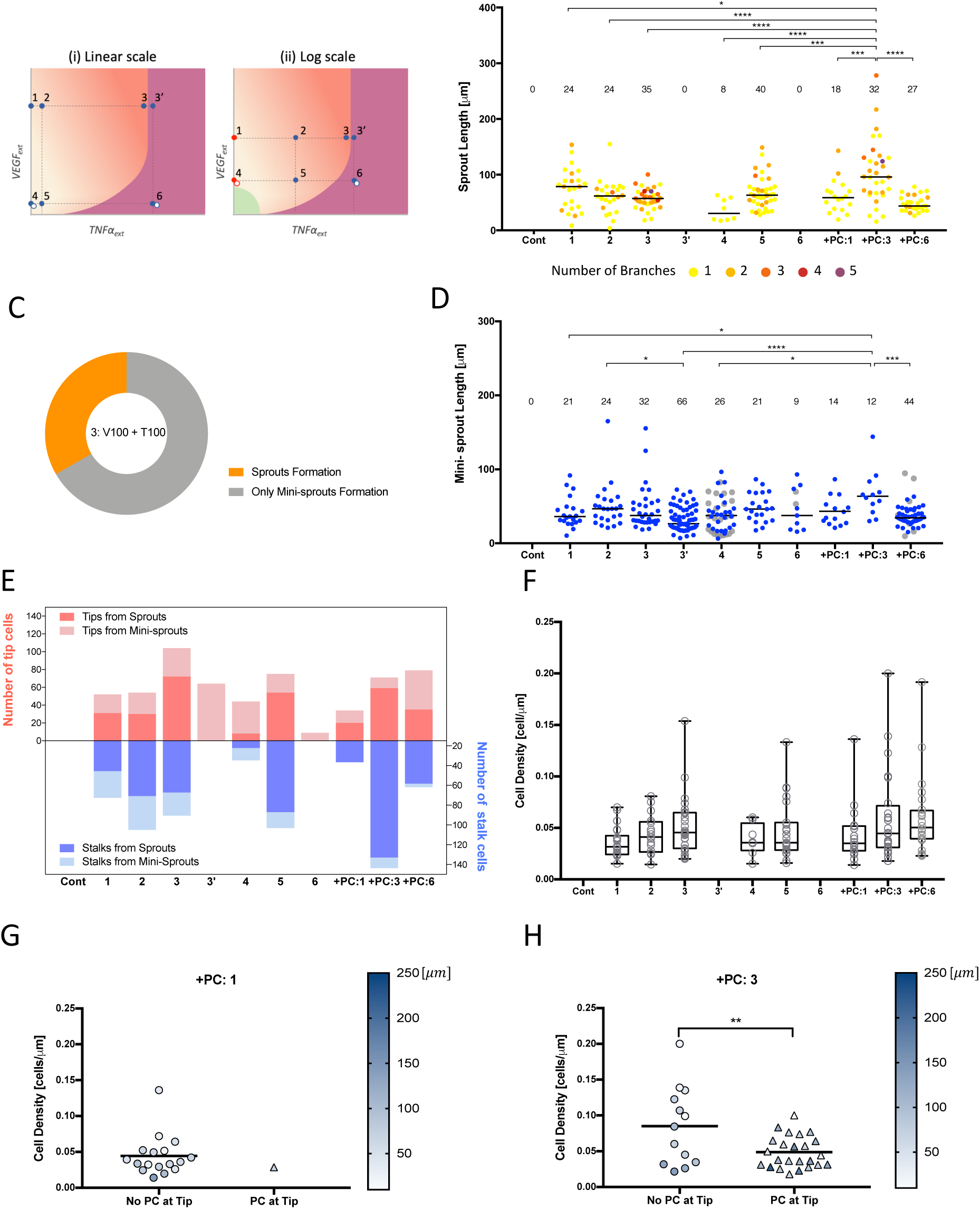
Pericytes binding substantially improve the stability of the Tip/Stalk differentiation process under inflammation. (A) Conceptual phase diagram of settled Tip-Stalk (T-S) decision phase (pale orange) and stalled Tip/Stalk-Tip/Stalk (T/S-T/S) decision phase (deep orange) displayed in linear scale (i) and logarithmic scale (ii). Point 1 and 4 are indicated with red dots in logarithmic scale due to the zero value of VEGF. Grey dots at point 4 and 6 represent the conditions presumably corresponding to those in Figure 2. The condition of point 3 and 3’ is the same but they are marked separately according to contradictory outcomes. (B) Quantification of lumenized sprout formation according to TNF*α* and VEGF concentrations specified in table 1. Color codes from 1 to 5 of sprouts indicate the number of branches. (C) Two possible and conflicting outcomes at point 3 from six independent experiments. The data from each experiment is presented in Figures S10. (D) Quantification of single cell-sized mini-sprouts formation according to TNF*α* and VEGF concentrations specified in table 1. Data from Figures 2E and F are displayed at 4, 6, +PC:6 with grey dots. (E) Quantification of the number of leading tip cells and following stalk cells from newly lumenized sprouts. Escalating TNF levels were initially pro-angiogenic but there was a complete abrogation of the formation of lumenized sprouts beyond a critical TNF level (point 3’ and point 6). This anti-angiogenic effect of TNF was completely rescued if the experiment was repeated in the presence of pericytes (+PC:3 and +PC:6). (F) Cell density of tubes which is calculated by dividing the number of cells consisting a sprout by its length. Increase of cell density along with TNF*α* level represented the stalk cell growth promoted by TNF*α*, resulting in thinker sprout formation. Cell density of +PC: 1 condition (G) and +PC: 3 condition (H) which was separated upon the presence of pericytes at the tip of sprouts. Color scale represents the length of sprout. Pericytes maintaining physically close contact with tip cells guided endothelial cells to form longer and thinner sprouts. Sprouts of +PC:6 were too short to differentiate pericytes at tip and stalk.

## Discussion

The effect of the local inflammation and the associated cytokines on the onset and progression of angiogenesis is still a matter of debate^5, 7-9^. In particular, it has been shown that while TNF can show strong anti-angiogenic effects^5, 7^, it can also induce the Tip cell fate through upregulation of a Notch ligand Jag-1, which might result in a pro-angiogenic function^8, 9, 27, 31^. Here, we demonstrate, through a combination of controlled experimentation with inducible angiogenesis in an engineered system and a computational model trained on the experimental data, that these apparently contradictory findings can be reconciled within a single framework. Furthermore, this framework accounts for diverse combinatorial effects of pro-angiogenic factors, such as VEGF, and pro-inflammatory cytokines, such as TNF, modulated by unanticipated juxtacrine influence of pericytes. The mathematical model capturing the details of the signaling networks involved in specification of the Tip and Stalk cells predicted the existence of the critical TNF concentrations, above which the interactions between Dll4, Jag1 and Notch can no longer induce differentiation between neighboring endothelial cells into distinct fates, yielding instead an intermediate or hybrid state. We also note that TNF, at high levels, may have an indirect effect on Notch signaling, by loosening cell-junctions^32-35^and thus diminishing juxtacrine Notch receptor engagement. Lack of cell differentiation above a critical TNF level, in turn, abrogates successful sprout induction, since if a presumptive Tip cell is not supported by neighboring proliferative Stalk cells, it cannot successfully migrate into the surrounding matrix without severing its contacts with the other cells. This observation is consistent with our findings of stunted single cell-sized ‘sprouts’ protruding from the parental vessel at TNF levels above the critical concentration. Likewise, if a Stalk fate induction is not supported by a neighboring Tip cell specification, the proliferative capacity inherent in the Stalk cell phenotype will be suppressed by the contact inhibition from neighboring cells within the endothelium due to the lack of Stalk cell separation from the rest of the monolayer. This is consistent with the apparent absence of Stalk cells at TNF concentrations above critical levels.

We found that pericyte coverage can rescue the inhibitory effect of TNF on angiogenesis, due to suppression of TNF signaling in endothelial cells. Although the VEGF signaling in these cells is also inhibited by the juxtacrine effects of pericytes, the influence of pericytes on TNF signaling is more consequential, due to the properties of the thresholds, separating the pro- and anti-angiogenic effects of TNF. At almost all VEGF levels, the values of these thresholds are predicted to be effectively independent of the VEGF concentration. Therefore, suppressing VEGF signaling will have little effect on the critical TNF concentrations, although VEGF can modulate the probability of induction of Tip and Stalk cells in a dose dependent fashion, if angiogenesis proceeds^15^.

The effects of pericytes on angiogenesis are predicted to strongly depend on the pericyte coverage of endothelial cells. This finding is important since pericyte coverage can vary across tissues, and between normal and cancerous vasculature, as well as be dynamic due to transient pericyte dissociation at the outset of angiogenesis^20, 36, 37^. For a very low pericyte coverage, the effect of TNF would be fully inhibitory, if this cytokine exceeded the critical level. On the other hand, a high, uniform pericytes coverage, affecting all endothelial cells can rescue the TNF-mediated suppression of angiogenesis. More interestingly, for an intermediate pericyte coverage levels, such that for many neighboring endothelial cells only one cell in two would be in contact with a pericyte, the pericyte-mediated rescue of the inhibitory TNF effect can dramatically increase, essentially rendering TNF strongly pro-angiogenic. This effect, as predicted by the model, would occur due to an enhanced differentiation of neighboring endothelial cells, guided by a higher TNF signaling in a cell that is not in contact with a pericyte and a lower signaling in a neighboring cell in contact with a pericyte. This asymmetry in signaling, coupled with the additional differentiation mediated by Notch signaling, can enhance the Tip/Stalk fate specification and promote the emergence and maturation of the nascent sprouts. This effect was consistent with the particularly pronounced growth and branching of the sprouts under conditions (high TNF levels) otherwise inhibitory to angiogenesis in the presence of pericytes. It also was so site that with the frequent observation of a pericyte at the tip areas of the particularly long sprouts, suggesting that the Tip/Stalk fate selection may be stabilized by a combination of a partial pericyte coverage and TNF, not only during the onset of sprout extension, but also during the sprout growth. Overall, our integrative analysis suggests an unexpected conclusion that angiogenesis can be particularly enhanced, leading to longer and more branched blood vessels, at relatively high ambient TNF levels in the presence of partial coverage by pericytes.

These findings are interesting to put into the context of the other commonly accepted views on the functions of pericytes within the exiting vascular beds. Low and intermediate pericyte coverage has been suggested to result in lower stability and higher leakage of blood vessels, which might be an indication of either pathologic conditions (e.g., in the context of growing tumors) or dynamically re-organizing vasculature. This view is consistent with our observations, suggesting that high pericyte coverage can down-modulate the effects of TNF and possibly other relevant signaling inputs, thus protecting the cells from the environmental stimuli, which may otherwise decrease the stability of the vessel. The destabilizing effect of low pericyte coverage can in part reflect a more variable effective sensitivity of endothelial cells to various extracellular stimuli, leading to more extensive angiogenesis, which might not however result in optimal functionality, unless pericyte coverage can be recovered through recruitment or differentiation of precursor cells.

*In vivo*, the local microenvironment can be highly dynamic. In particular, successful angiogenesis might lead to a gradual restoration of appropriate oxygen tension and also be accompanied by a progressive inflammation resolution. As these conditions evolve, the biochemical milieu may change along with dynamic alterations of inputs from mural cells, such as pericytes, raising the question of how the resulting morphogenesis of vascular beds might be affected. Our analysis provides a useful framework that can help start analyzing the complex multi-factorial control of this morphogenetic process critical in a variety of developmental and physiological settings. More generally, it can also provide an insight into how heterotypic cell interactions can also modulate Notch signaling in other contexts, regulating cellular differentiation outcomes.

## References

1. Adams, R.H. & Alitalo, K. Molecular regulation of angiogenesis and lymphangiogenesis. Nat Rev Mol Cell Biol 8, 464–478 (2007).

2. Potente, M., Gerhardt, H. & Carmeliet, P. Basic and therapeutic aspects of angiogenesis. Cell 146, 873–887 (2011).

3. Albini, A. & Sporn, M.B. The tumour microenvironment as a target for chemoprevention. Nat Rev Cancer 7, 139–147 (2007).

4. Eming, S.A., Krieg, T. & Davidson, J.M. Inflammation in wound repair: molecular and cellular mechanisms. J Invest Dermatol 127, 514–525 (2007).

5. Sainson, R.C. et al. TNF primes endothelial cells for angiogenic sprouting by inducing a tip cell phenotype. Blood 111, 4997–5007 (2008).

6. Madge, L.A. & Pober, J.S. TNF signaling in vascular endothelial cells. Exp Mol Pathol 70, 317–325 (2001).

7. Frater-Schroder, M., Risau, W., Hallmann, R., Gautschi, P. & Bohlen, P. Tumor necrosis factor type alpha, a potent inhibitor of endothelial cell growth in vitro, is angiogenic in vivo. Proc Natl Acad Sci U S A 84, 5277–5281 (1987).

8. Boareto, M., Jolly, M.K., Ben-Jacob, E. & Onuchic, J.N. Jagged mediates differences in normal and tumor angiogenesis by affecting tip-stalk fate decision. Proc Natl Acad Sci U S A 112, E3836–3844 (2015).

9. Benedito, R. et al. The Notch Ligands Dll4 and Jagged1 Have Opposing Effects on Angiogenesis. Cell 137, 1124–1135 (2009).

10. Chen, M.B. et al. On-chip human microvasculature assay for visualization and quantification of tumor cell extravasation dynamics. Nat Protoc 12, 865–880 (2017).

11. Nguyen, D.H. et al. Biomimetic model to reconstitute angiogenic sprouting morphogenesis in vitro. Proc Natl Acad Sci U S A 110, 6712–6717 (2013).

12. Bayless, K.J., Kwak, H.I. & Su, S.C. Investigating endothelial invasion and sprouting behavior in three-dimensional collagen matrices. Nat Protoc 4, 1888–1898 (2009).

13. Jakobsson, L. et al. Endothelial cells dynamically compete for the tip cell position during angiogenic sprouting. Nat Cell Biol 12, 943–953 (2010).

14. Bentley, K. et al. The role of differential VE-cadherin dynamics in cell rearrangement during angiogenesis. Nat Cell Biol 16, 309–321 (2014).

15. Noren, D.P. et al. Endothelial cells decode VEGF-mediated Ca2+ signaling patterns to produce distinct functional responses. Sci Signal 9, ra20 (2016).

16. Eilken, H.M. & Adams, R.H. Dynamics of endothelial cell behavior in sprouting angiogenesis. Curr Opin Cell Biol 22, 617–625 (2010).

17. Hoang, M.V., Whelan, M.C. & Senger, D.R. Rho activity critically and selectively regulates endothelial cell organization during angiogenesis. Proc Natl Acad Sci U S A 101, 1874–1879 (2004).

18. Bryan, B.A. et al. RhoA/ROCK signaling is essential for multiple aspects of VEGF-mediated angiogenesis. Faseb J 24, 3186–3195 (2010).

19. Gerhardt, H. & Betsholtz, C. Endothelial-pericyte interactions in angiogenesis. Cell Tissue Res 314, 15–23 (2003).

20. Bergers, G. & Song, S. The role of pericytes in blood-vessel formation and maintenance. Neuro Oncol 7, 452–464 (2005).

21. Maier, C.L., Shepherd, B.R., Yi, T. & Pober, J.S. Explant outgrowth, propagation and characterization of human pericytes. Microcirculation 17, 367–380 (2010).

22. Clewley, R. Hybrid models and biological model reduction with PyDSTool. PLoS Comput Biol 8, e1002628 (2012).

23. Herland, A. et al. Distinct Contributions of Astrocytes and Pericytes to Neuroinflammation Identified in a 3D Human Blood-Brain Barrier on a Chip. PLoS One 11, e0150360 (2016).

24. Campisi, M. et al. 3D self-organized microvascular model of the human blood-brain barrier with endothelial cells, pericytes and astrocytes. Biomaterials 180, 117–129 (2018).

25. Kim, J. et al. Engineering of a Biomimetic Pericyte-Covered 3D Microvascular Network. PLoS One 10, e0133880 (2015).

26. Lee, E. et al. A 3D in vitro pericyte-supported microvessel model: visualisation and quantitative characterisation of multistep angiogenesis. J Mater Chem B 6, 1085–1094 (2018).

27. Johnston, D.A., Dong, B. & Hughes, C.C. TNF induction of jagged-1 in endothelial cells is NFkappaB-dependent. Gene 435, 36–44 (2009).

28. Thurston, G. & Kitajewski, J. VEGF and Delta-Notch: interacting signalling pathways in tumour angiogenesis. Br J Cancer 99, 1204–1209 (2008).

29. Selvam, S., Kumar, T. & Fruttiger, M. Retinal vasculature development in health and disease. Prog Retin Eye Res 63, 1–19 (2018).

30. Shaya, O. & Sprinzak, D. From Notch signaling to fine-grained patterning: Modeling meets experiments. Curr Opin Genet Dev 21, 732–739 (2011).

31. Petrovic, J. et al. Ligand-dependent Notch signaling strength orchestrates lateral induction and lateral inhibition in the developing inner ear. Development 141, 2313– 2324 (2014).

32. Dewi, B.E., Takasaki, T. & Kurane, I. In vitro assessment of human endothelial cell permeability: effects of inflammatory cytokines and dengue virus infection. J Virol Methods 121, 171–180 (2004).

33. Aveleira, C.A., Lin, C.M., Abcouwer, S.F., Ambrosio, A.F. & Antonetti, D.A. TNF-alpha signals through PKCzeta/NF-kappaB to alter the tight junction complex and increase retinal endothelial cell permeability. Diabetes 59, 2872–2882 (2010).

34. Shaya, O. et al. Cell-Cell Contact Area Affects Notch Signaling and Notch-Dependent Patterning. Dev Cell 40, 505–511 e506 (2017).

35. Clark, P.R., Kim, R.K., Pober, J.S. & Kluger, M.S. Tumor necrosis factor disrupts claudin-5 endothelial tight junction barriers in two distinct NF-kappaB-dependent phases. PLoS One 10, e0120075 (2015).

36. Zhang, L. et al. Presence of retinal pericyte-reactive autoantibodies in diabetic retinopathy patients. Sci Rep 6, 20341 (2016).

37. Huang, F.J. et al. Pericyte deficiencies lead to aberrant tumor vascularizaton in the brain of the NG2 null mouse. Dev Biol 344, 1035–1046 (2010).

