## Supplemental Figures for "Pericytes enable effective angiogenesis in the presence of pro-inflammatory signals"

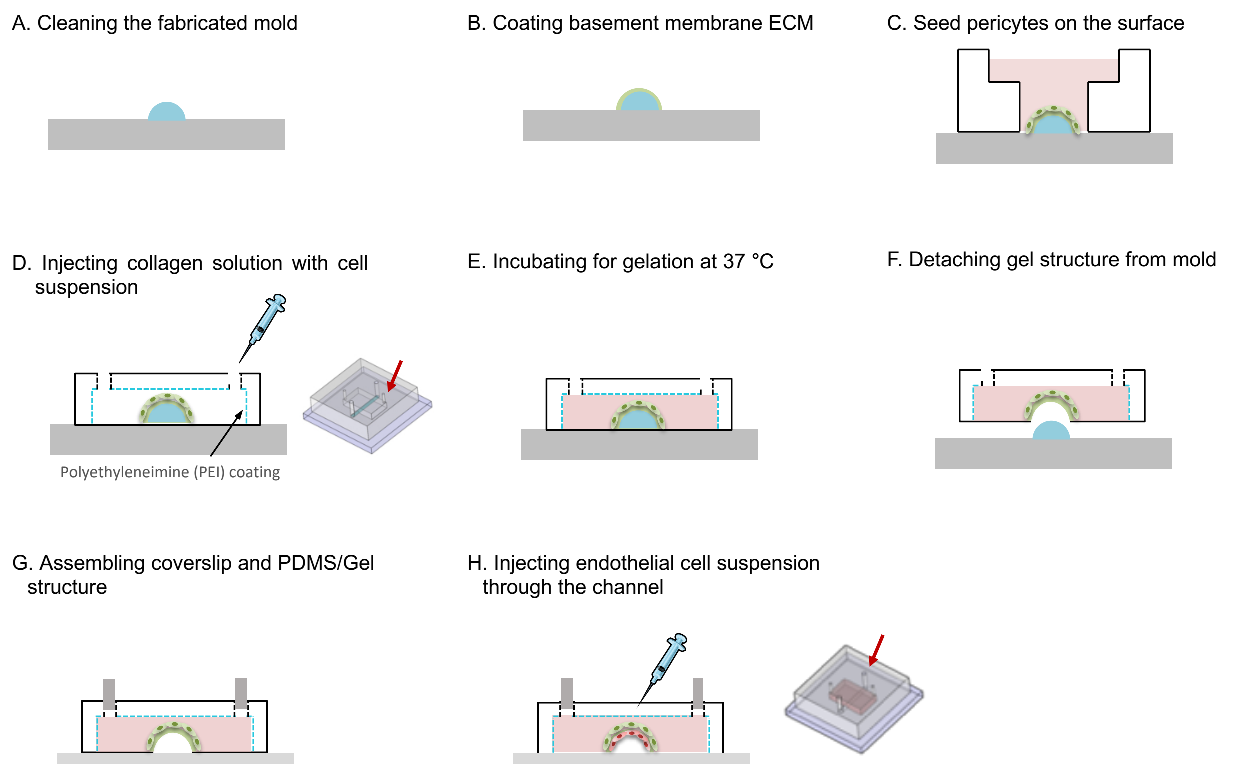


**Figure S1. Fabrication process of 3D vessel on a chip.** (A) A single line mold (D: 200 -250 um, L: 10 mm) which has a semicircular cross-section was deposited on a petri dish. (B)The mold was covered completely by 4% collagen (Corning) and the remnant was aspirated gently by pipetting. EDC (1-Ethyl-3-(3-dimethylaminopropyl)-carbodiimide)/NHS (N-hydroxysulfoxuccinimide) solution in ethanol (10mg/m) was placed on the mold from the side and it was left at room temperature for 1min for inducing collagen cross-linking, washed with DI water and the solution on the mold was completely aspirated. The same process was repeated three times and the mold was washed with DI water three times at the end. (C) PDMS reservoir was put on the mold and 30 µl of pericytes suspension ($6 \times 10^5$ cells/ml) was placed in the reservoir. The pericytes seeded mold was incubated in 37 ℃ for 5 hours. For 3D vessel chip without pericytes, this process was skipped. (D) The PDMS chamber and cover slip was pre-treated with 0.1% (v/v) poly(ethylenimine) (PEI) for 30 min, 1% glutaraldehyde for 1hr. After taking off the reservoir,

PDMS chamber was put on the mold. Pre-mixed collagen solution (5mg/ml, Type 1 collagen, BD) according to manufacturer's protocol was injected through the hole (indicated with red arrow) and (E) collagen polymerization was induced on ice for 30 min and in 37 ℃ for 1.5 hours. (F) The single line mold was carefully peeled off, (G) then the bottom side of PDMS chamber containing collagen gel was sealed to a glass coverslip. (H) Endothelial cell suspension ($1 \times 10^7$cells /ml) was injected through the inlet hole (indicated with red arrow), then the chip was flipped upside down and incubated in 37 ℃ for 1 hour to allow cells to attach on the lumen. Fresh medium was injected through the inlet of the cannel

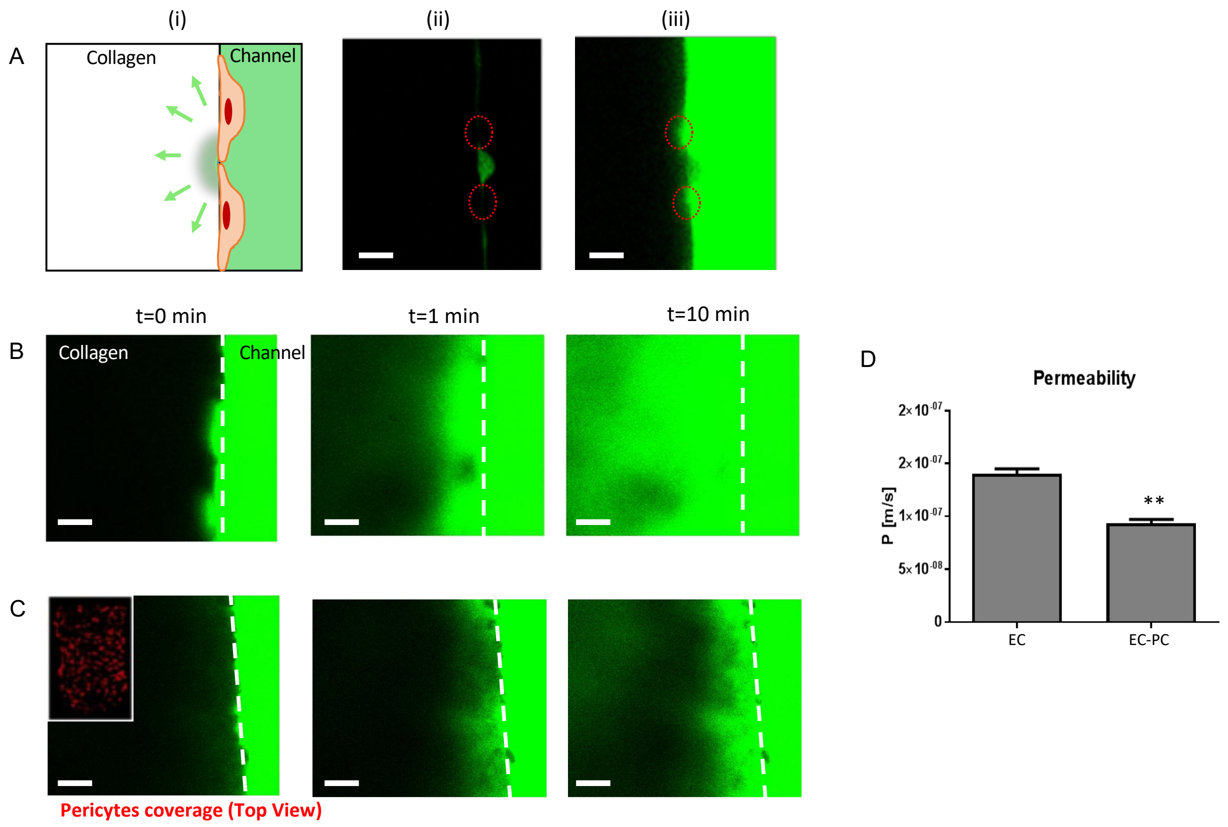


**Figure S2. Permeability evaluation with FITC-Dextran (70kDa).** (A) Cell culture medium containing FITC-dextran was injected through the channel (i) then FITC-Dextran diffused primarily through cell-cell junction (ii &iii). Endothelial cell was pre-labeled with DiO (green). FITC-Dextran diffusion through bare endothelium (B) and through endothelium covered with pericytes (C). Pericytes were pre-labeled with DiI (red). Scale bar, 100 $\mu m$. (D) Permeability evaluation with FITC-Dextran (70 kDa) shows the increase of vessel stabilization by pericytes, resulting in the decrease of permeability. **P<0.01.

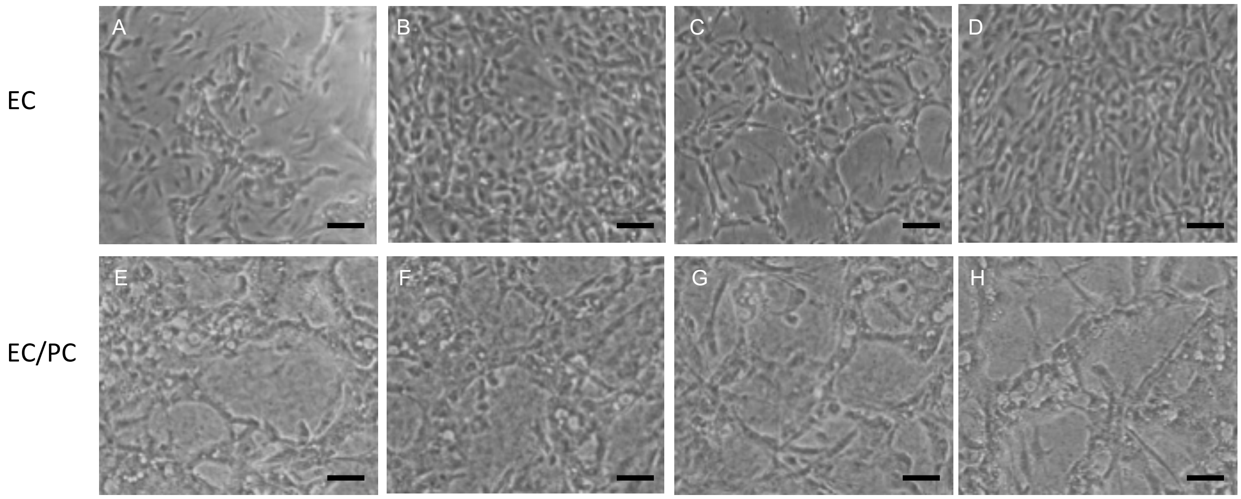


**Figure S3. Vasculogenesis on sandwich culture.** Formation of tube-like structure with normal medium (A), with 100 ng/ml of TNFα (B), with 100 ng/ml of VEGF (C), with 100 ng/ml of TNFα and 100ng/ml of VEGF (D) on EC monolayer, formation of tube-like structure with normal medium (E), with 100ng/ml of TNFα (F), with 100 ng/ml of VEGF (G), with 100 ng/ml of TNFα and 100 ng/ml of VEGF (H) on EC/PC mixed monolayer. Scale bar, 100 $\mu m$

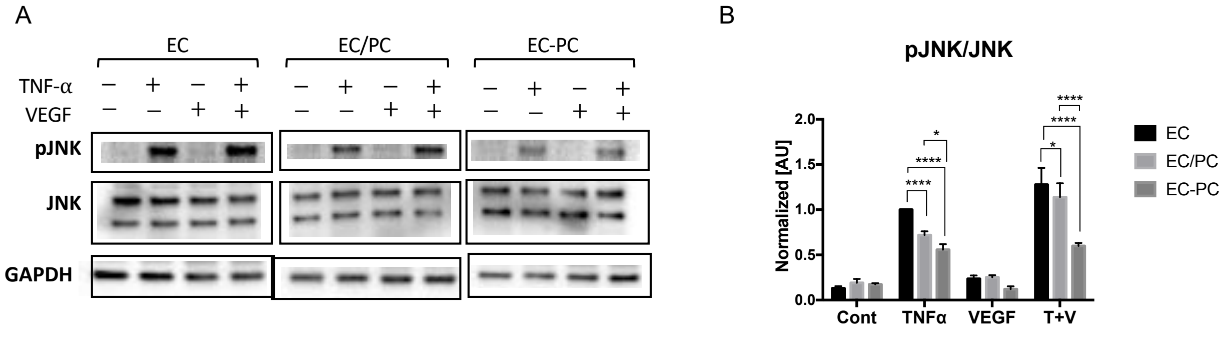


**Figure S4. JNK activity regulated by direct binding of pericytes.** (A) Western blot images of phospho-JNK and JNK, (B) quantified data from the images.

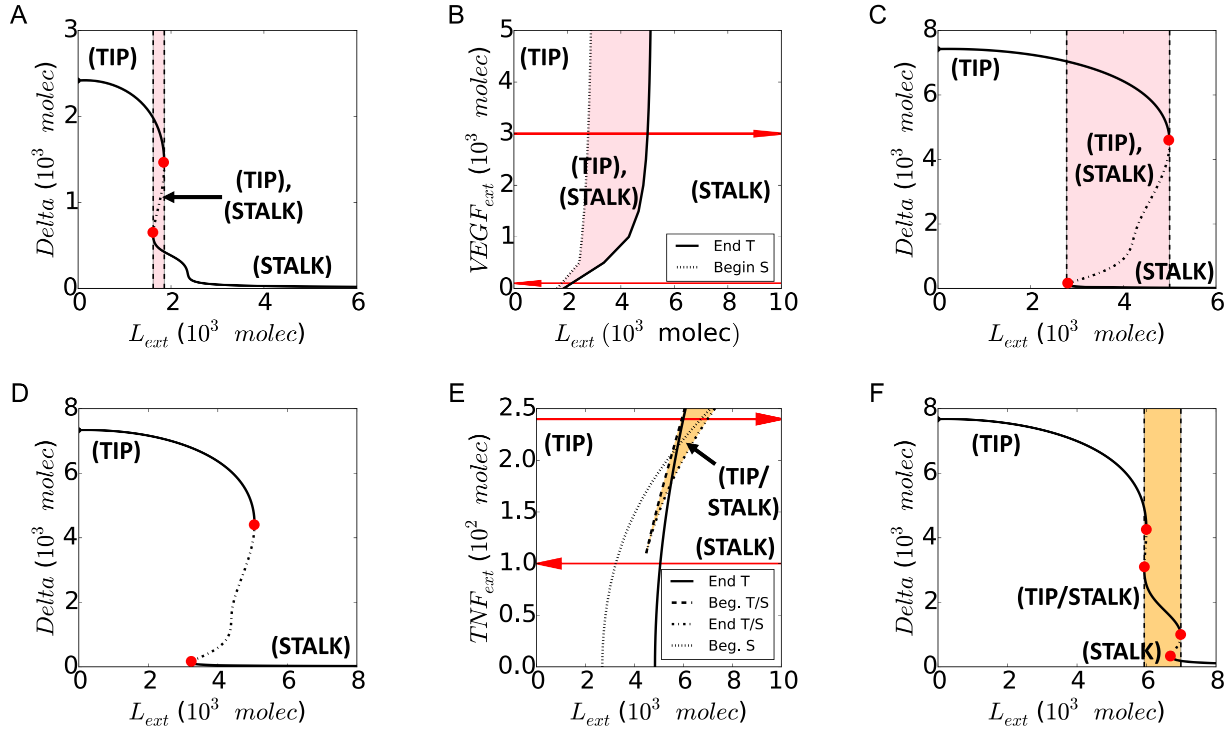


**Figure S5 Effect of external signals on cell fate determination in the single-cell model.** (A) Bifurcation diagram of intra-cellular Delta with the external Notch ligand ($L_{ext} = D_{ext} + J_{ext}$) as control parameter and low exposure to external VEGF signal ($VEGF_{ext} = 0$ molecules, see red arrow in panel (B)). Continuous lines indicate stable fixed points, while dotted lines indicate unstable fixed points. (B) Phenotype phase diagram of the single cell exposed to external Notch ligands (x-axis, $L_{ext}$) and external VEGF signal (y-axis, $VEGF_{ext}$). Tip (T) and Stalk (S) phenotypes coexist for a strong VEGF signal (pink-shaded area indicates bi-stability). (C) Same as (A) for a high VEGF external signal ($VEGF_{ext} = 3000$ molecules, see top red arrow in panel (B)). (D) Bifurcation diagram of intracellular Delta with the external Notch ligands ($L_{ext}$) as control parameter and a low exposure to external TNF signal ($TNF_{ext} = 0$ molecules, see bottom red arrow in panel (E)). (E) Phenotype phase diagram of the single cell exposed to external Notch ligands (x-axis, $L_{ext}$) and TNF signal (y-axis, $TNF_{ext}$). TNF signal introduces a hybrid Tip/Stalk phenotype (T/S) (orange-shaded area). (F) Same as (D) for high TNF external signal ($TNF_{ext}$ = 240 molecules, see top red arrow in panel (E)). In panels (A), (B) and (C), $TNF_{ext} = 0$ molecules, while in panels (D), (E) and (F) $VEGF_{ext} = 2000$ molecules. Bifurcation diagrams of all other variables corresponding to (A)-(C)-(D)-(F) are presented in Fig. S6-S7

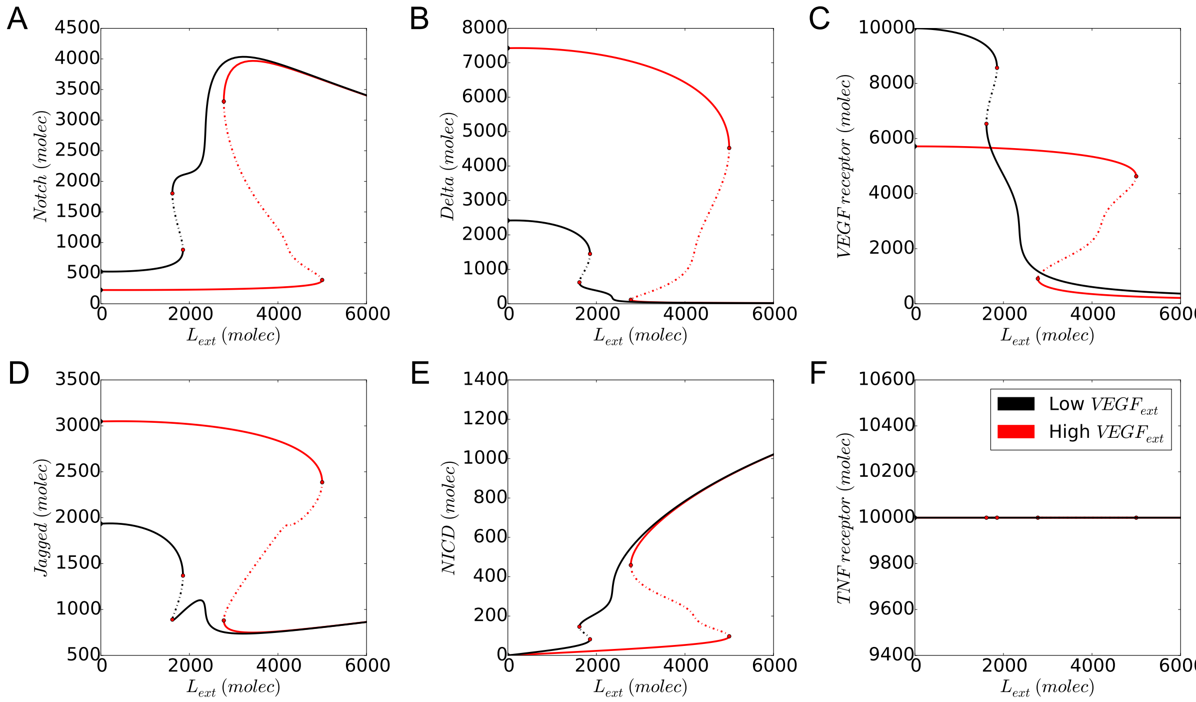


**Figure S6. Bifurcation diagram of the model's variables for low and high exposure to VEGF signaling.** Bifurcation diagram of (A) Notch, (B) Delta (Dll4), (C) VEGF receptor, (D) Jagged (Jag1), (E) NICD and (F) TNF receptor with external Notch ligands ($L_{ext} = D_{ext} + J_{ext}$) as control parameter. Black curves indicate low exposure to external VEGF signaling (as of Fig. S5A) while red lines indicate high exposure (as of Fig. S5C). $TNF_{ext} = 0$ molecules in this simulation.

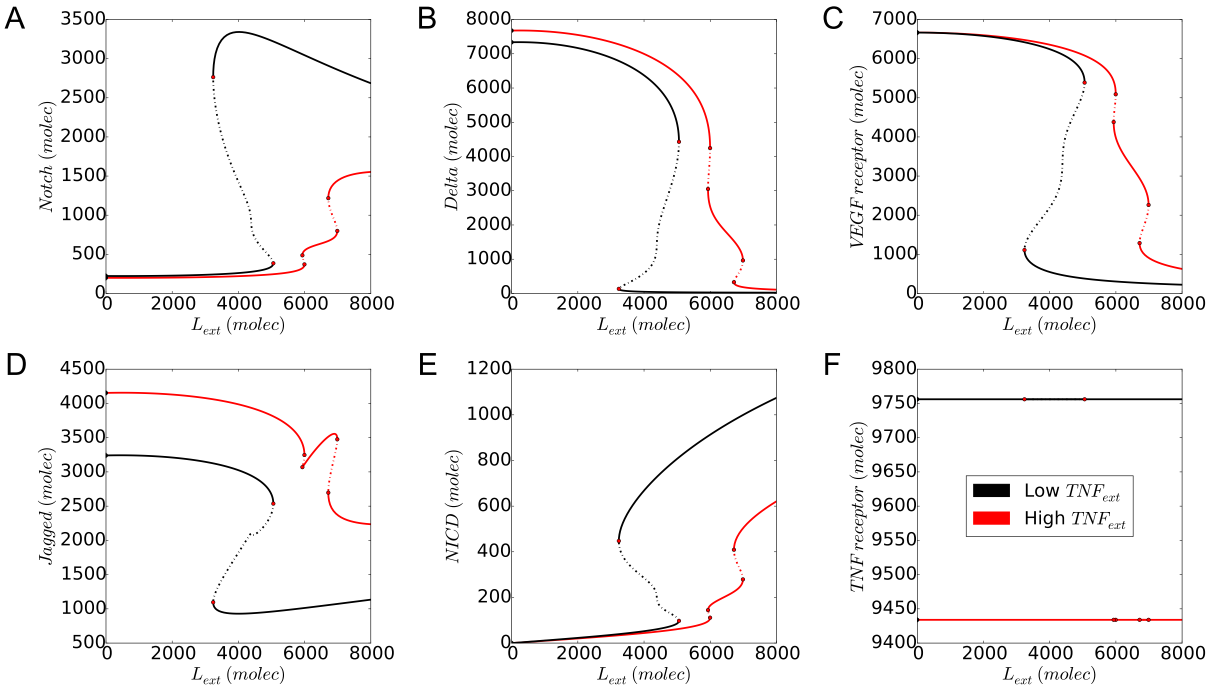


**Figure S7 Bifurcation diagram of the model's variables for low and high exposure to TNF signaling.** Bifurcation diagram of (A) Notch, (B) Delta (Dll4), (C) VEGF receptor, (D) Jagged (Jag1), (E) NICD and (F) TNF receptor with external Notch ligands ($L_{ext} = D_{ext} + J_{ext}$) as control parameter. Black curves indicate low exposure to external TNF signaling (as of Fig. S5d) while red lines depict high exposure to external TNF signaling (as of Fig. S5F). $VEGF_{ext} = 2000$ molecules in this simulation.

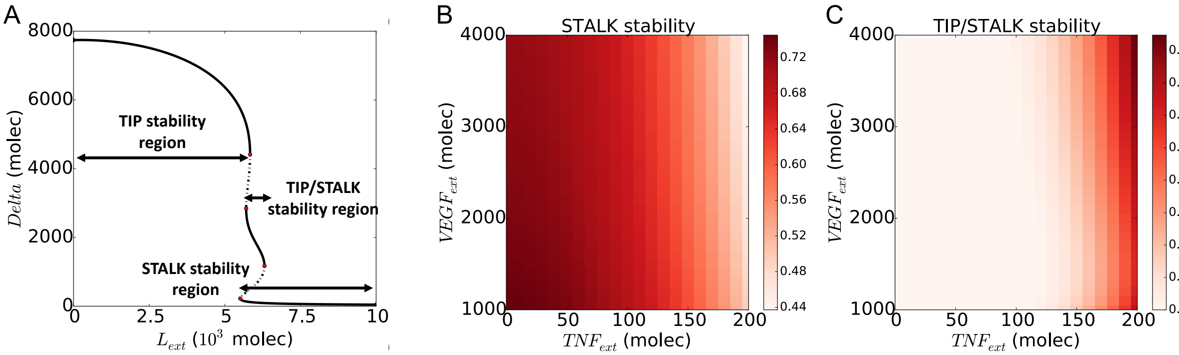


**Figure S8 Stabilization/destabilization of each state by VEGF and TNF signaling.** (A) Intracellular Delta (Dll4) as a function of external Notch ligands, the stability of STALK (B) and TIP/STALK (C) depending on the level of $VEGF_{ext}$ and $TNF_{ext}$ in each region of (A). The stability of a certain cell state corresponds to the range of the parameter $L_{ext}$ on the x-axis that allows that state, divided by the total range of the x-axis.

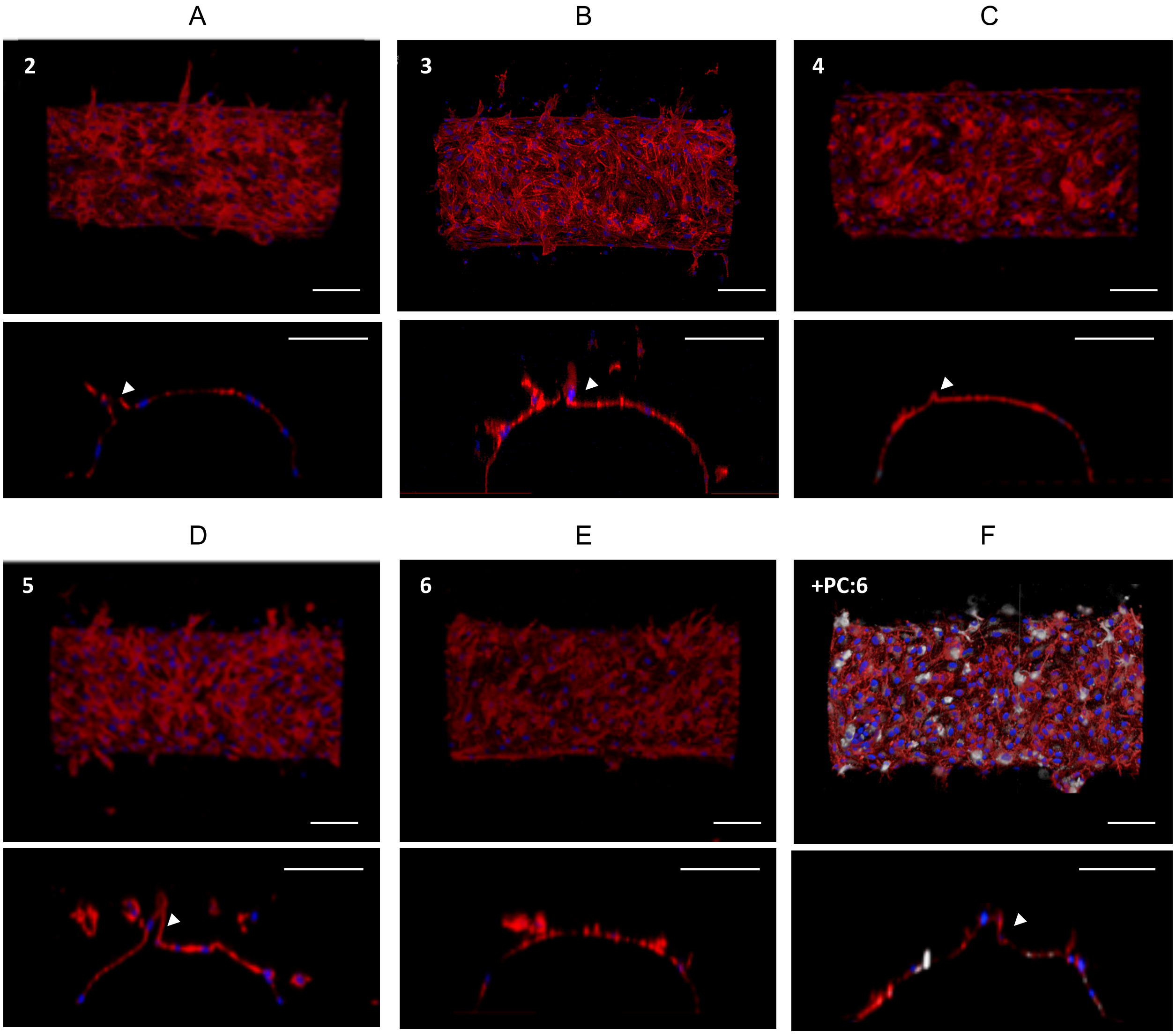
A
B
C
2
3
4
D
E
F
5
6
+PC:6

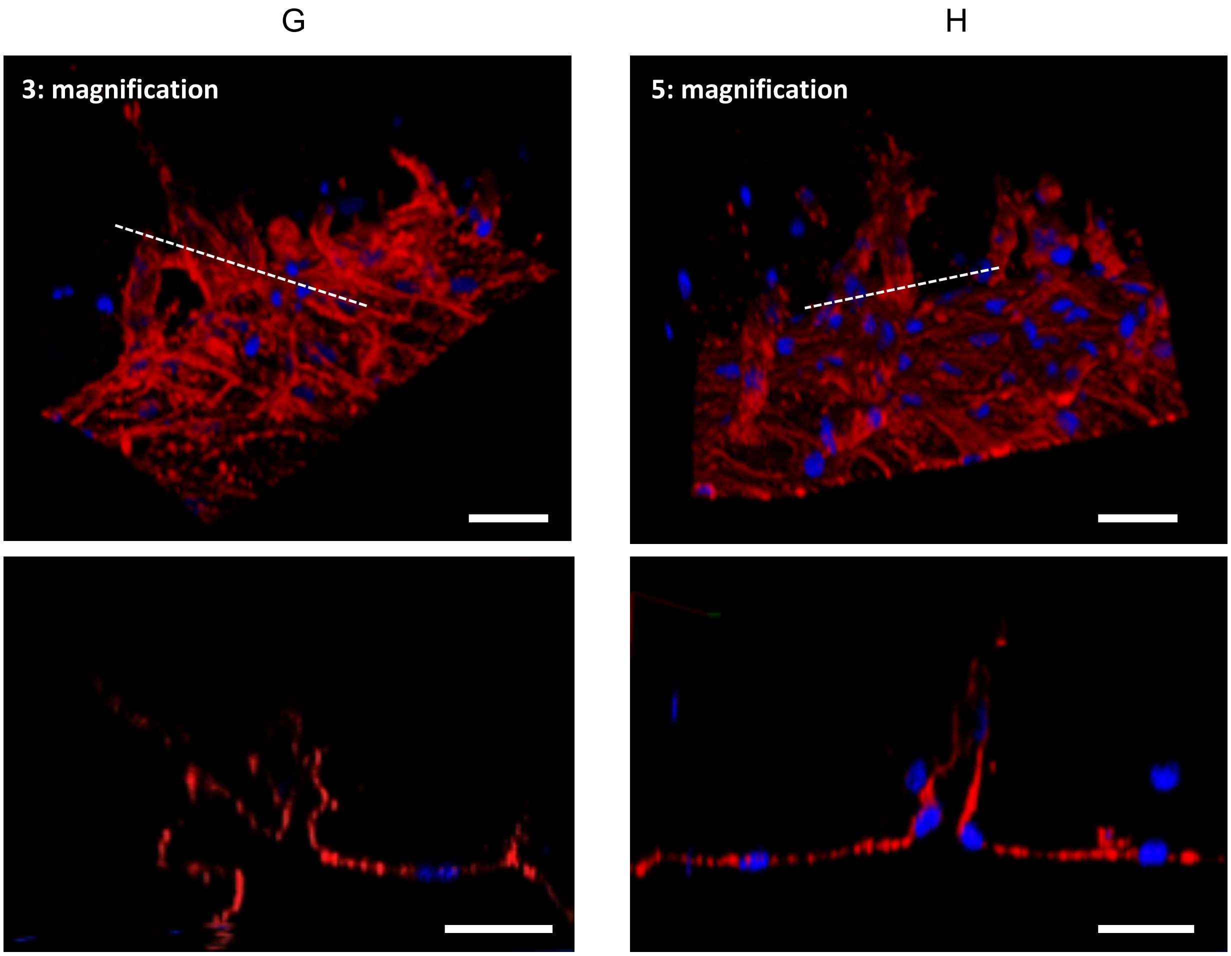


**Figure S9. Angiogenesis in 3D vessel on a chip in response to gradient of VEGF and local TNF$\boldsymbol{\alpha}$.** Confocal images of sprouting and tube formation in response to the condition of 2 (A), 3 (B), 4 (C), 5 (D), 6 (E), and 6 with pericytes (F) indicated in Table 1. Scale bar, 100 $\mu m$. The magnified cross-section images of condition 3 (G) and 5 (H) show highly branched and irregular tube formation and elongated single tube formation, respectively. Scale bar, 25 $\mu m$. Actin filaments of endothelial cells and pericytes were stained with phalloidin (red) and pericytes were pre-labeled with DiD (white).

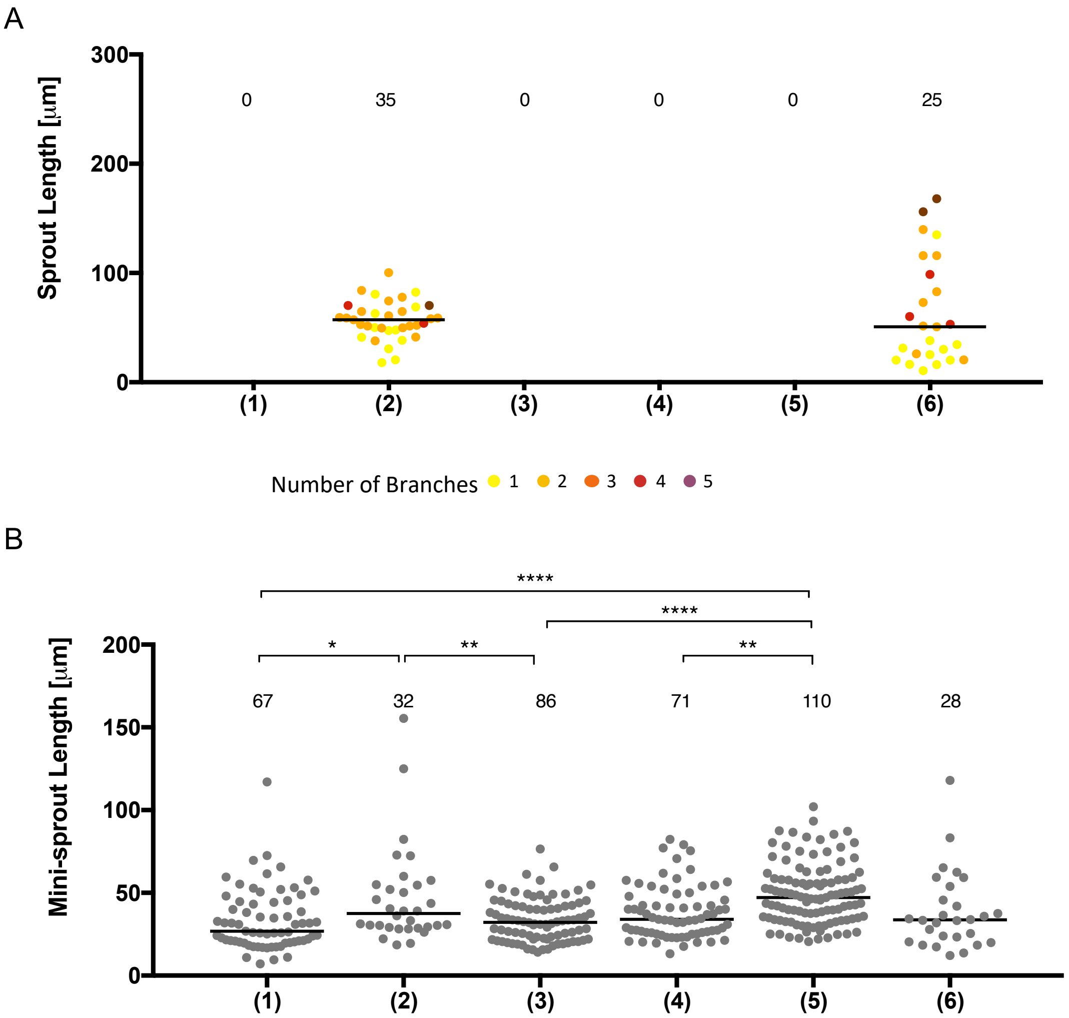


**Figure S10. Angiogenic response at condition 3 in table 1.** (A) Quantification of lumenized sprout formation. Color codes from 1 to 5 of sprouts indicate the number of branches. (B) Quantification of single cell-sized mini-sprouts formation.
