## Supplemental Information for "Pericytes enable effective angiogenesis in the presence of pro-inflammatory signals"

### Mathematical model of the Notch-VEGF-TNF circuit

The model of the Notch-VEGF-TNF circuit comprehends six variables: Notch receptor ($N$), Delta ($D$) and Jagged ($J$) ligands, Notch IntraCellular Domain (NICD, $I$), VEGF receptor ($V_R$) and TNF receptor ($T_R$). Here, Delta and Jagged correspond to Dll4 and Jag1, the ligand subtypes considered in the experimental system. The temporal dynamics of the intra-cellular levels of the six variables were described via a system of ordinary differential equations:

$$\frac{dN}{dt} = N_0 H^{S+}(I, \lambda_{I,N}) - N[(k_C D + k_T D_{ext}) + (k_C J + k_T J_{ext})] - \gamma N$$

$$\frac{dD}{dt} = D_0 p_D H^{S-}(I, \lambda_{I,D}) H^{S+}(k_T V_R V_{ext}/\gamma_S, \lambda_{V,D}) - D[k_C N + k_T N_{ext}] - \gamma D$$

$$\frac{dJ}{dt} = J_0 p_J H^{S+}(I, \lambda_{I,J}) H^{S+}(k_T T_R T_{ext}/\gamma_S, \lambda_{T,J}) - J[k_C N + k_T N_{ext}] - \gamma J$$

$$\frac{dI}{dt} = k_T N[D_{ext} + J_{ext}] - \gamma_S I$$

$$\frac{dV_R}{dt} = V_{R0} H^{S-}(I, \lambda_{I,V_R}) - k_T V_R V_{ext} - \gamma V_R$$

$$\frac{dT_R}{dt} = T_{R0} - k_T T_R T_{ext} - \gamma T_R$$

In the model, all variables are expressed in number of molecules. $N_0$, $D_0$, $J_0$, $V_{R0}$, $V_{T0}$ are the basal cellular production rate constants of $N$, $D$, $J$, $V_R$ and $T_R$. Since only the molecular copy number is considered in the model, these rate constants account for both transcription and translation. All quantities degrade at the same rate $\gamma$, while $I$ degrades at a faster rate $\gamma_S$. The transcriptional regulation that $I$ exerts on any other component $X$ ($N$, $D$, $J$ or $V_R$) was modeled via shifted Hill functions $H^{S+/-}$ defined as:

$$H^{S+/-}(I, I_0, n_{I,X}, \lambda_{I,X}) = \frac{1}{1+\left(\frac{I}{I_0}\right)^{n_{I,X}}} + \lambda_{I,X}\frac{\left(\frac{I}{I_0}\right)^{n_{I,X}}}{1+\left(\frac{I}{I_0}\right)^{n_{I,X}}}$$

where +/- stands for activation $(N, J)$ or inhibition $(D, V_R)$ and $I_0$ is a half-maximal level of $I$. The Hill coefficient $n_{I,X}$ relates to the steepness of the regulation with respect to $I$ and $\lambda_{I,X}$ is a fold-change in the production rate of $X$ due to $I$ ($\lambda_{I,X} < 1$ for inhibition, $\lambda_{I,X} > 1$ for activation)(2). The binding of $V_R$ to external VEGF molecules ($V_{ext}$) leads to activation of $D$. This interactions was described via the shifted Hill function $H^{S+}(k_T V_R V_{ext}/\gamma_S, \lambda_{V,D})$, where the amount of active intra-cellular VEGF signaling $k_T V_R V_{ext}/\gamma_S$ is explicitly derived in the following paragraph. Similarly, $J$ is activated upon binding of $T_R$ to external TNF molecules ($T_{ext}$). This interaction was modeled via the shifted Hill function $H^{S+}(k_T T_R T_{ext}/\gamma_S, \lambda_{T,J})$, where $k_T T_R T_{ext}/\gamma_S$ is the level of active intra-cellular TNF signaling (see next paragraph). $N_{ext}$, $D_{ext}$ and $J_{ext}$ represent a constant level of external Notch receptor and ligands in the single cell model or the level of Notch receptor and ligands in the neighboring cell for the 2-cell model. In the latter case, each cell is described by its own set of variables for the Notch, VEGF and TNF pathways. $V_{ext}$ and $T_{ext}$ are the levels of external VEGF and TNF ligands, and are assumed to be constant for both the single cell and the 2-cell model. $k_C$ and $k_T$ are the cis- and trans- receptor-ligand binding rate constants. In the model, only $N$ can bind with ligands produced in the same cell (cis-interaction), leading to degradation of the complex. $V_R$ and $T_R$ can only bind to external molecules (trans-interaction). $p_D$ and $p_J$ are numerical factors that decrease the production rates of $D$ and $J$ in presence of pericytes. In this case, the production rate constants of D and J are simply rescaled as:

$$D_0 \rightarrow p_D D_0$$

$$J_0 \rightarrow p_J J_0$$

Since pericytes have an inhibitory effect on D and J, $p_D$ and $p_J$ can be varied between 0 (complete shutdown of D, J production) and 1 (control case with no effect of pericytes).

**Determination of active VEGF/TNF intracellular signaling**

The level of active signaling $A$ (valid for both VEGF or TNF) is described by the equation:

$$\frac{dA}{dt} = k_T A_R A_{ext} - \gamma_S A$$

where $k_T$ is the binding rate of the receptor $A_R$ with the external signaling molecule $A_{ext}$ and $\gamma_S$ is a degradation rate constant. By assuming fast equilibration $\frac{dA}{dt} = 0$, the steady state level of active signaling becomes:

$$A = \frac{k_T A_R A_{ext}}{\gamma_S}$$

therefore, $\frac{k_T V_R V_{ext}}{\gamma_S}$ and $\frac{k_T T_R T_{ext}}{\gamma_S}$ represent the level of active VEGF and TNF signaling in the cell, respectively.

**Parameters estimation**

The parameters of the Notch-VEGF axis were taken from the original model of Boareto et al(1) without modifications. In the original model, the values of the production rates of $N$, $D$, $J$ and $V_R$ ensured an order of magnitude of $\sim 10000$ molecules at cell surface(3). The Hill function parameters were estimated from seminal experimental data. The Supplementary Information of Boareto et al(1) provides to the interested reader the detailed derivation and discussion of the pre-existing parameters. The degradation rate of $T_R$ is equal to the degradation rate of $N$, $D$, $J$ and $V_R$ ($\gamma = 0.1$, typical of protein degradation(4)). The production rate $T_{R0}$ equals the production rate of VEGF ($1000$ mRNA $h^{-1}$) to ensure a similar receptor level. Similarly, the ligand-receptor binding rate for TNF equals the trans-interaction rate of $N$ and $V_R$ ($k_T = 5\ 10^{-4}\ h^{-1}$). The fold-change in the production of $J$ due to TNF ($\lambda_{T,J} = 5$) was inferred from the experimental data presented in this paper (Fig .3G), while the Hill coefficient was assumed to be $n_{T,J} = 2$. The threshold level of active

TNF signal ($T_{0,J} = 400$ molecules) was chosen to enable a tangible effect on $J$, given the baseline level of $T_R$ in the model. The fold-changes $p_D$ and $p_J$ were estimated from the experimental data presented in this paper (Figs. 3G & H). All parameters are listed in Table S1.

| **Parameter group** | **Parameter** | **Value** | **Dimensions** |
|---|---|---|---|
| Degradation | $\gamma, \gamma_S$ | $0.1, 0.5$ | $h^{-1}$ |
| Production | $N_0, D_0, J_0, V_{R0}, T_{R0}$ | $1200, 1000, 800, 1000, 1000$ | mRNA $h^{-1}$ |
| Hill coefficient | $n_{I,N}, n_{I,D}, n_{I,J}, n_{I,V}, n_{V,D}, n_{T,J}$ | $2, 2, 5, 2, 2, 2$ | Dimensionless |
| Hill fold-change | $\lambda_{I,N}, \lambda_{I,D}, \lambda_{I,J}, \lambda_{I,V}, \lambda_{V,D}, \lambda_{T,J}$ | $2, 0, 2, 0, 2, 5$ | Dimensionless |
| Hill threshold | $I_{0,N}, I_{0,D}, I_{0,J}, I_{0,V}, V_{0,D}, T_{0,J}$ | $200, 200, 200, 200, 200, 400$ | molecules |
| Binding rate constants | $k_T, k_C$ | $5\ 10^{-4}, 2.5\ 10^{-5}$ | $h^{-1}$ |
| Pericytes | $p_D, p_J$ | $0.6, 0.6$ | Dimensionless |

*Table S1. Parameters of the Notch-VEGF-TNF circuit.*

**Simulation details**

The bifurcation and phase diagrams of the single cell model were calculated using the Python numerical library PyDSTool(5). The code for the calculations of the 2-cell model was developed by the Federico Bocci, and it is available upon request. The system of ODEs that describes the 2-cell system was solved using an explicit Eulerian numerical integration scheme, and the level of active VEGF signaling in the two cells were taken upon equilibration.

**References**


1. Boareto M, Kumar M, Ben-jacob E, Onuchic JN. Jagged mediates differences in normal and tumor angiogenesis by affecting tip-stalk fate decision. 2015;
2. Lu M, Kumar M, Levine H, Onuchic JN, Ben-jacob E. MicroRNA-based regulation of epithelial-hybrid mesencymal fate determination. Proc Natl Acad Sci. 2013;110(45):18144–18149.
3. Boareto M, Jolly MK, Lu M, Onuchic JN, Clementi C, Ben-Jacob E. Jagged--Delta asymmetry in Notch signaling can give rise to a Sender/Receiver hybrid phenotype. Proc Natl Acad Sci. 2015;112(5):E402--E409.
4. Eden E, Geva-zatorsky N, Issaeva I, Cohen A, Dekel E, Cohen L, et al. Proteome Half-Life Dynamics in Living Human Cells. Science. 2011;331(6018):764–8.
5. Clewley R. Hybrid models and biological model reduction with PyDSTool. PLoS Comput Biol. 2012;8(8):e1002628.
